# Development of FERM domain protein-protein interaction inhibitors for MSN and CD44 as a potential therapeutic strategy for Alzheimer’s disease

**DOI:** 10.1101/2023.05.22.541727

**Authors:** Yuhong Du, William J. Bradshaw, Tina M. Leisner, Joel K. Annor-Gyamfi, Kun Qian, Frances M. Bashore, Arunima Sikdar, Felix O. Nwogbo, Andrey A. Ivanov, Stephen V. Frye, Opher Gileadi, Paul E. Brennan, Allan I. Levey, the Emory-Sage-SGC TREAT-AD Center, Alison D. Axtman, Kenneth H. Pearce, Haian Fu, Vittorio L. Katis

## Abstract

Recent genome-wide association studies have revealed genetic risk factors for Alzheimer’s disease (AD) that are exclusively expressed in microglia within the brain. A proteomics approach identified moesin (MSN), a FERM (four-point-one ezrin radixin moesin) domain protein, and the receptor CD44 as hub proteins found within a co-expression module strongly linked to AD clinical and pathological traits as well as microglia. The FERM domain of MSN interacts with the phospholipid PIP_2_ and the cytoplasmic tails of receptors such as CD44. This study explored the feasibility of developing protein-protein interaction inhibitors that target the MSN–CD44 interaction. Structural and mutational analyses revealed that the FERM domain of MSN binds to CD44 by incorporating a beta strand within the F3 lobe. Phage-display studies identified an allosteric site located close to the PIP_2_ binding site in the FERM domain that affects CD44 binding within the F3 lobe. These findings support a model in which PIP_2_ binding to the FERM domain stimulates receptor tail binding through an allosteric mechanism that causes the F3 lobe to adopt an open conformation permissive for binding. High-throughput screening of a chemical library identified two compounds that disrupt the MSN–CD44 interaction, and one compound series was further optimized for biochemical activity, specificity, and solubility. The results suggest that the FERM domain holds potential as a drug development target. The small molecule preliminary leads generated from the study could serve as a foundation for additional medicinal chemistry effort with the goal of controlling microglial activity in AD by modifying the MSN–CD44 interaction.

## Introduction

Alzheimer’s disease (AD) is a progressive neurodegenerative disorder that results in dementia, affecting over 50 million people worldwide (1). AD pathology is characterized by the build-up within the brain of amyloid beta-containing senile plaques and the formation of neurofibrillary tangles composed of hyperphosphorylated tau. Despite efforts to target amyloid beta and tau for disease-modifying treatments, phase III trials have demonstrated limited success in slowing cognitive decline (2, 3). As our understanding of the complex pathophysiology of AD remains incomplete, there is a need to explore the role of other potential proteins in the disease and to evaluate their therapeutic feasibility. In recent years, inflammatory signalling within microglia has received increasing attention for its role in AD pathology (4), with a number of proteins linked to microglial signalling identified as potential drug targets (5). Chronic activation of microglia can lead to the production of toxic cytokines and chemokines, which cause neuronal damage. Suppressing microglial-mediated inflammation may be an important strategy in the treatment of AD.

To gain a more comprehensive understanding of AD biology, Johnson *et al*. (2020) utilized an unbiased proteomics approach to examine tissue from more than 400 human brains. Co-expression network analysis identified a module (M4) that was enriched with AD risk genes and proteins expressed in both astrocytes and microglia (6, 7). Within this module, two hub proteins, moesin (MSN) and CD44, were found to have strongly correlated expression with cognition, amyloid plaque and neurofibrillary tangle burden. Given the module’s enrichment in genes associated with AD risk, these hub proteins are considered as causal drivers in disease progression (8). Protein levels of MSN and CD44 are elevated in AD brains, with CD44 levels enriched in AD patient cerebral spinal fluid (6, 8). MSN, which is predominantly produced in microglial cells in the brain, was often observed surrounding amyloid beta plaques in AD patients and the 5xFAD mouse model (7). CD44, on the other hand, is produced in both neuronal and glial cells, where it is involved in neuro-inflammation (9, 10). Loss of function studies highlight a role for CD44 in amyloid-beta induced neurotoxicity (11).

The ERM (ezrin radixin moesin) family of proteins, to which MSN belongs, is involved in linking the actin cytoskeleton to the plasma membrane (12, 13). This allows ERM proteins to regulate cytoskeletal rearrangements that govern various cellular functions, including migration, motility, cell shape, and signalling. ERM proteins are localized to specialized phosphatidylinositol 4,5-bisphosphate (PIP_2_)- enriched regions within the membrane (14, 15), where their N-terminal FERM (four-point-one ezrin radixin moesin) domain binds to PIP_2_ and the cytoplasmic C-terminal tails of receptor proteins like CD44. The C-terminal domain of ERM proteins blocks the binding of CD44 to the FERM domain when in an inactive conformation (16, 17). However, phosphorylation of this inhibitory domain results in its dissociation from the FERM domain, enabling it to bind to CD44 (18) (as depicted in Fig. 1A). The FERM domain is made up of three subdomains (F1, F2, and F3) that form a ‘three-leaf clover’ topology. Structural investigations of mouse radixin indicate that CD44 binds to subdomain F3 (19). CD44, a cell surface glycoprotein, is a receptor for hyaluronic acid, a crucial component of the extracellular matrix in different organs, including the brain. By associating with ERM proteins, the binding of CD44 to hyaluronic acid through its extracellular domain is believed to promote cell adhesion and migration (20).

**Figure 1:**
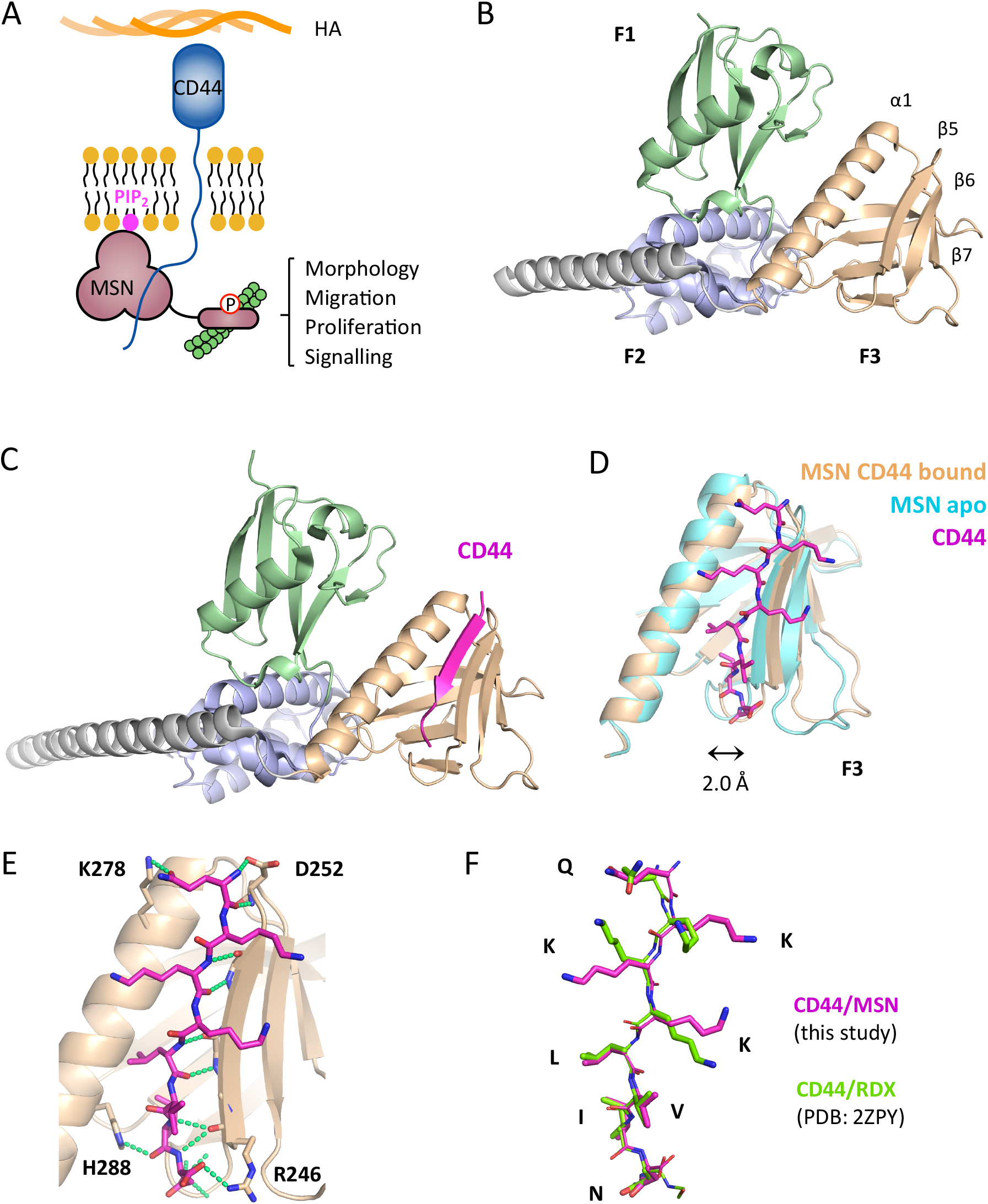
Crystal structure of MSN-FERM, either ligand free or bound to CD44 cytoplasmic tail. (**A**) Model showing MSN activation through binding the C-terminal cytoplasmic tail of the CD44 receptor and PIP_2_. (**B**) Crystal structure of ligand-free FERM domain of human MSN (PDB: 6TXQ). The three FERM subdomains of MSN, shown as ribbons, are coloured in green (F1), metallic blue (F2) and wheat colour (F3), while the long C-terminal alpha helix is coloured in grey. (**C**) Crystal structure of human MSN bound to a short synthetic CD44 peptide (678-QKKKLVIN-685). CD44 (magenta, ribbon) binds within the F3 domain, forming part of the beta-sheet. (**D**) CD44 within the F3 lobe (magenta, stick model) is accommodated by a displacement of the beta sheet away from the alpha helix by 2.0 Å, measured from H288 and R246 carbonyl carbon atoms. MSN bound to CD44 is shown in wheat colour, while unbound MSN is shown in cyan. (**E**) Interactions between MSN and CD44, showing hydrogen bonding (green dashed lines). The majority of bonding is backbone-backbone, although a few MSN sidechain residue interactions (H288, D252, R246) were identified. The MSN-K278 interaction is most likely an artefact of crystal packing. (**F**) Overlay of the FERM domain-bound CD44 to either MSN (this work) or radixin (PDB: 2ZPY).

The above evidence suggests that both MSN and CD44 are involved in the inflammatory response observed in the brains of AD patients. Although it is unclear if MSN plays a detrimental or protective function in AD pathogenesis (7), inhibition of the interaction between these two proteins is proposed to reduce neuronal damage by hyperactive microglia. In this study we explore the tractability of developing small molecule inhibitors that disrupt the interaction of the FERM domain of MSN with CD44. Co-crystallography revealed the FERM domain binds to the cytoplasmic tail of CD44 in a similar fashion to that seen previously with radixin (19). A simple and robust biochemical assay based on time-resolved fluorescence resonance energy transfer (TR-FRET) was developed to measure the MSN–CD44 interaction. High throughput screening identified two compounds that could displace this interaction, both *in vitro* and *in cellulo*. Further optimisation improved biochemical activity, specificity and solubility of one of these compounds. To investigate whether MSN–CD44 protein-protein interaction (PPI) inhibitors could be obtained that bind at allosteric sites on the FERM domain, we employed a peptide phage-display library. In conclusion, this study demonstrates that the FERM domain is a viable target for drug development, and further refinement of preliminary leads could provide valuable insight into the role of MSN and CD44 in AD.

## Results

### High resolution structure of MSN FERM domain

To assist our efforts in obtaining MSN–CD44 PPI inhibitors, we determined the crystal structure of the FERM domain of human MSN to a resolution of 1.73 Å (Fig. 1B, PDB: 6TXQ) (Table S1). We compared our structure to a previously published lower resolution MSN FERM-domain crystal structure (1E5W; 2.7 Å) (21). Despite sharing the same overall topology, there were notable differences between the two structures (Fig. S1A). In our structure, the F3 lobe was significantly moved away from the F1 subdomain, which was rotated by 12° along the long alpha helix (Fig. S1B). While the F1 and F2 subdomains were very similar between both structures, significant changes were observed within the F3 lobe (Fig. S1C). Our structure had the extended helix (α1) and beta sheet (β5–β7) in a “closed” conformation, which could not accommodate a beta strand from a receptor tail such as that of CD44. In contrast, the previous MSN-FERM structure (1E5W) had a more “open” F3 subdomain with the beta sheet displaced from the helix by 2.5 Å (Fig. S1C). The variable positions of the subdomains observed in the two MSN FERM structures suggest they have a high degree of independent movement within the FERM domain.

### Structure of MSN FERM domain bound to CD44

We have successfully determined a crystal structure of the FERM domain of MSN in complex with a short peptide that includes the ERM binding region of CD44’s cytoplasmic tail, solved to 2.2 Å (Fig. 1C; PDB:6TXS) (Table S1). Within the F3 lobe of MSN, CD44 forms a beta strand that sits adjacent to the extended helix, contributing to the β5–β7 anti-parallel beta-sheet. This type of interaction was also seen in a previous study where a CD44 peptide was bound to radixin (19), an ERM protein that is closely related to MSN (2ZPY). Comparison of the CD44-bound and ligand free MSN structures revealed that peptide binding led to a significant displacement of the β5–β7 beta-sheet from the alpha helix (2.0 Å), as shown in Fig. 1D. The interaction between CD44 and MSN is mainly through backbone-backbone hydrogen bonds with the MSN β5 strand. Additionally, the CD44 peptide backbone can form hydrogen bonds with the H288 sidechain within the extended alpha helix and salt bridges to the R246 and D252 sidechains within the β5 strand (Fig. 1E). Interestingly, the R246 and D252 interactions were not observed in the radixin-CD44 structure (19), although these residues are conserved between both ERM proteins. The CD44-bound structures of both ERM proteins were found to have closely aligned peptide backbones, although the three lysine residues within the CD44 peptide were spread out in our structure, forming a large hydrophobic surface (Fig. 1F). While the electron density of the lysine sidechains from the radixin-CD44 structure was considerably weaker, our structure provided clear density with better B-factors for these sidechains (2ZPY: 60-80 Å^2^; 6TXS: 40-50 Å^2^). When bound to MSN, the 8-residue CD44 peptide was found to have a total buried surface area of 1392 Å^2^.

### A MSN TR-FRET assay reveals residues important for CD44 binding

A time-resolved fluorescence resonance energy transfer (TR-FRET) assay was established to measure the binding of CD44 peptide to MSN. The assay generated a specific FRET signal from purified 6His-tagged MSN bound to terbium and an FITC-conjugated CD44 peptide (Fig. 2A). The specificity of the TR-FRET signal was confirmed by titration with increasing concentrations of unlabelled CD44 peptide (Fig. 2B). Site-directed mutagenesis was used to validate and disrupt the CD44 interaction observed in the structure. However, due to the lack of MSN side-chain interactions to the peptide, we focussed our efforts on H288, a conserved residue among both ERM proteins and more distantly related FERM domain proteins, such as 4.1B and FRMD6. H288, which forms a hydrogen bond to the carbonyl of CD44 residue I684, was mutated to alanine. In addition, we mutated S249, L281 and M286 to arginine, a bulky charged residue that we predicted would cause steric hindrance with CD44 binding. While S249R and M286R showed no effect (not shown), H288A and L281R mutations resulted in weak TR-FRET signals compared to wild-type MSN (Fig. 2D) and required a 10-fold higher concentration to inhibit the wild-type MSN–CD44 interaction in a competition assay (Fig. 2E). The mutants showed minimal changes in melting temperature in a thermal shift assay (Fig. 2F) and crystal structures of MSN H288A and L281R mutant FERM domains confirmed that they were correctly folded (Fig. S2) (Table S1).

**Figure 2:**
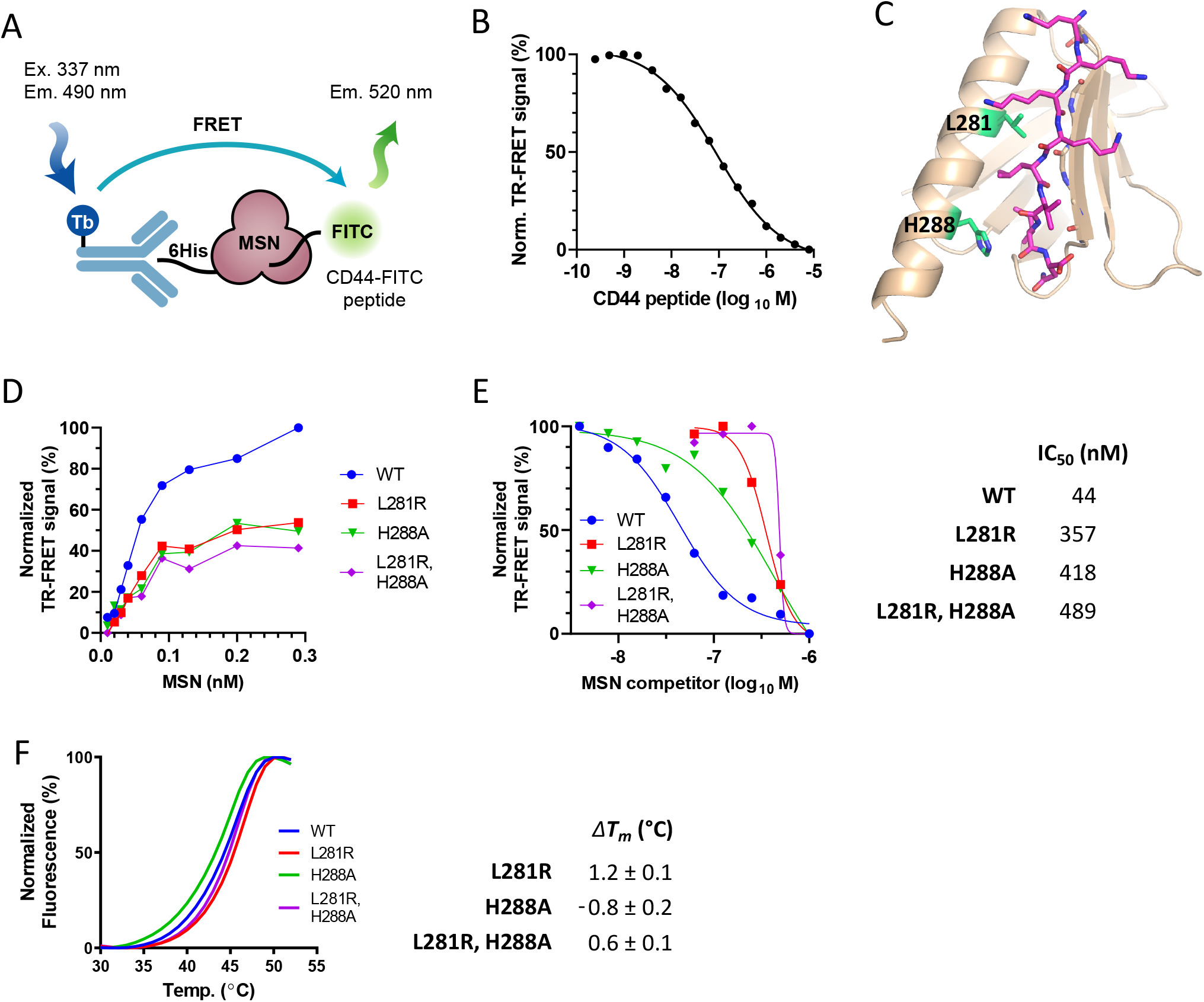
Structure-guided mutagenesis reveals MSN residues involved in CD44 binding. (**A**) Schematic representation of the *in vitro* TR-FRET assay for MSN–CD44 interaction using purified components. Signal generated from Tb-linked anti-6HIS antibody bound to MSN is transferred to the FITC-conjugated CD44 peptide. (**B**) Inhibition of the TR-FRET signal using unlabelled CD44 peptide as a competitor (IC_50_ = 66 nM). (**C**) Sidechains of mutated MSN residues (L281 and H288) within the MSN– CD44 binding pocket are shown (green). (**D**) Comparison of MSN–CD44 TR-FRET response using varying levels of 6His-tagged WT or MSN mutant protein (0.01-0.29 nM). CD44 peptide concentration was kept constant (8 nM) (**E**) Competition of mutant MSN protein in MSN–CD44 TR-FRET assay. Dose response of untagged WT or mutant MSN protein in TR-FRET assay containing wild-type 6His-MSN (2 nM) and CD44 peptide (8 nM). IC_50_ values of competitor proteins are shown on the right (n=3). (**F**) Thermal shift assay, showing melting temperature (*T*_*m*_) curves of WT and mutant proteins (left panel). The *ΔT*_*m*_, compared to WT protein, is shown in the right panel (n=3; ±SD).

### Phage display identifies peptides that bind to the MSN FERM domain at two distinct sites

To investigate possible allosteric binding pockets on the FERM domain of MSN that could affect CD44 interaction, we performed phage display screening using a library that contained a randomized 10-amino acid sequence. We screened the library both with and without the CD44 peptide and tracked each round of selection by monitoring colony formation units and phage ELISA. After examining the sequences of the clones that were identified as positive by ELISA, we discovered two peptides, C3P-pd and C3S1-pd, which were selected in the presence and absence of the CD44 peptide, respectively (Fig. 3A).

**Figure 3:**
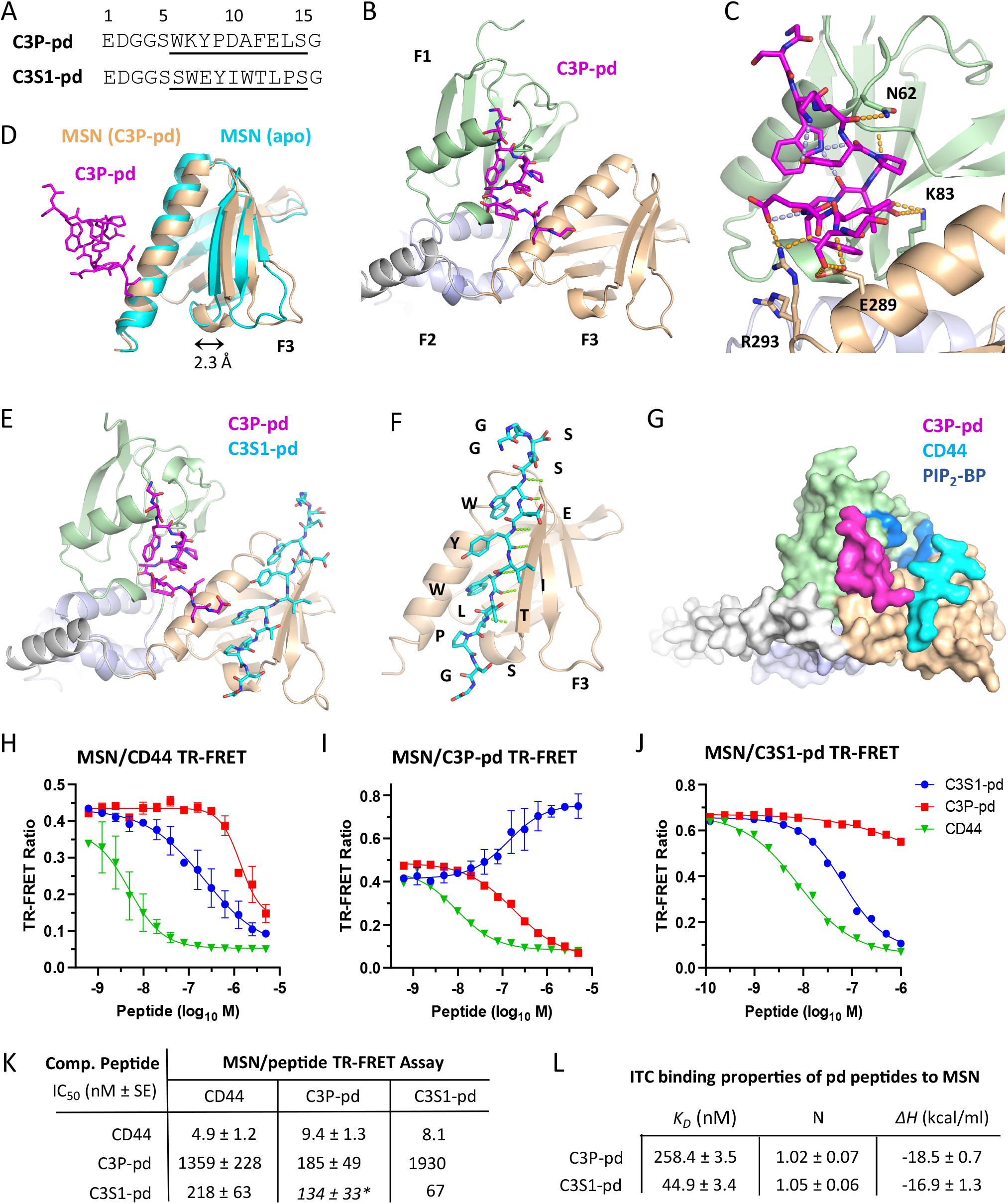
Phage display screening identifies peptides that bind MSN at distinct sites. (**A**) Sequences of phage display peptides. Underlined region corresponds to the variable region within the phage-derived sequence. (**B**) Crystal structure of FERM domain of MSN bound to the C3P-pd peptide. FERM subdomains are displayed as in **1B**. C3P-pd peptide (magenta) binds in a pocket between the F1 and F3 subdomains. (**C**) Close-up view of C3P-pd peptide bound to MSN. Peptide intramolecular H-bonds are shown (purple dashed lines). Sidechains of MSN residues interacting with the peptide are displayed, together with their bonding to the peptide (gold dashed lines). (**D**) Superimposed F3 subdomains of MSN from apo (cyan) and C3P-pd bound (light yellow) structures. C3P-pd peptide binding causes a 2.3 Å movement of the beta sheet away from the alpha helix in the MSN F3 lobe, measured from H288 and R246 carbonyl carbon atoms. (**E**) Crystal structure of FERM domain of MSN bound to both C3P-pd and C3S1-pd peptides. (**F**) Close-up of C3S1-pd binding to F3 lobe of MSN. H-bond contacts are shown (green dashed lines). (**G**) Space-filling model of MSN FERM domain, showing C3P-pd (magenta) and CD44 (cyan) binding relative to proposed PIP_2_ binding pocket (PIP_2_-BP; blue positively-charged surface). (**H, I, J**) MSN TR-FRET inhibition assays, with acceptor fluorophore conjugated to either (**H**) CD44, (**I**) C3P-pd, or (**J**) C3S1-pd peptides. Unlabelled peptides were used as competitors (C3S1-pd, blue circle; C3P-pd, red square; CD44, green triangle). (**K**) IC_50_ values derived from **H, I**, and **J**. Asterisk denotes stimulatory rather than inhibitory values (**L**) Binding properties MSN association to C3P-pd and C3S1-pd peptides, measured by ITC. See Figure S3 for thermograms and corresponding fitted curves. *K*_*D*_=dissociation constant, N=stoichiometry, *ΔH*=enthalpy.

A crystal structure of C3P-pd bound to MSN was determined (1.85 Å), which revealed the peptide binds to a crevice situated between the F1 and F3 subdomains, adjacent to the proposed PIP_2_ binding pocket (Fig. 3B and 3G) (Table S1). The peptide adopted an unusual compact structure that formed a plug between the two subdomains, supported by multiple internal hydrogen bonds, including the one formed between the indole nitrogen of W6 and two carbonyl groups within the peptide backbone. The peptide interacted with both subdomains via several hydrogen bonds (Fig. 3C). Specifically, within the F1 subdomain, C3P-pd interacted with the N62 and K83 sidechains. Within the F3 subdomain, the peptide interacted with the E289 and R293 sidechains from the extended alpha helix. Remarkably, binding of C3P-pd to MSN resulted in a 2.3 Å shift of the β5–β7 beta-sheet from the alpha helix in the F3 lobe, when compared to our MSN structure that lacked any ligands. This caused the subdomain to adopt an “open” conformation similar to that observed in the MSN–CD44 structure. This is surprising, since C3P-pd does not contact the beta sheet, suggesting an allosteric effect on the CD44 binding pocket.

Attempts to co-crystallize C3S1-pd and MSN were only successful in the presence of C3P-pd. The ternary complex structure, solved to 1.52 Å, contained both peptides bound, with C3S1-pd situated within the CD44 binding pocket (Fig. 3E) (Table S1). C3S1-pd, like CD44, forms a beta strand within the F3 lobe that binds in an antiparallel fashion to the β5–β7 beta sheet, displacing it from the alpha helix by 2.4 Å (not shown). Unlike CD44, C3S1-pd exclusively forms hydrogen bonds with the beta sheet (Fig. 3F). Although in close proximity, the C3P-pd and C3S1-pd peptides do not contact each other. The structure also contained a third bound peptide (not shown) on the two-fold axis between copies of the asymmetric unit. This peptide is likely to have been either C3P-pd or C3S1-pd, but we were unable to identify it due to poor sidechain density, so it was modelled as penta-alanylglycine.

We further characterized the phage-display peptides for their ability to displace CD44 in the TR-FRET assay. C3S1-pd effectively competed with CD44, demonstrating an IC_50_ of around 220 nM (Fig. 3H and 3K). In contrast, C3P-pd could only compete at high concentrations (IC_50_ =1.4 µM). This is not surprising, since C3P-pd was identified from selections carried out in the presence of the CD44 peptide. In addition, C3P-pd would not be predicted to sterically hinder CD44 when comparing superimposed structures of C3P-pd and CD44-bound MSN (Fig. 3F). TR-FRET assays were also developed using Cy5-conjugated phage display peptides (Fig. 3I and 3J). Interestingly, CD44 was effective at displacing C3P-pd (IC_50_=9.4 nM), suggesting inhibition through an allosteric mechanism. Surprisingly, C3S1-pd increased the FRET signal generated from MSN and C3P-pd-Cy5, showing an EC_50_ of around 130 nM. This suggests that C3S1-pd can allosterically stimulate binding of C3P-pd. However, the converse was not true, as C3P-pd had little effect on C3S1 binding (Fig. 3J and 3K). Binding properties of the phage display peptides were measured by isothermal titration calorimetry (ITC). Both peptides bound to MSN in a 1:1 ratio, with C3P-pd and C3S1-pd having dissociation constants (*K*_*D*_) of around 260 and 45 nM, respectively (Fig. 3L and S3).

### uHTS screening for MSN–CD44 PPI inhibitors

We miniaturized and optimized the MSN–CD44 TR-FRET assay into a 1536-well ultra-high throughput screening (uHTS) format to conduct screening against a chemical diversity compound library with the goal of obtaining small molecule inhibitors capable of disrupting the MSN–CD44 interaction. Details of assay optimization, miniaturization and screening will be published separately as an assay development and screening technical resource for scientific community. A chemical library composed of 138,000 compounds was screened, giving a primary hit rate of 0.2% (Fig. 4A). After removing known PAINS (pan-assay interference compounds) (22) from the primary list, we cherry-picked the remaining hits from the library stock and confirmed their activity through dose-response in the TR-FRET assay. The compounds were then re-ordered and confirmed once more by TR-FRET dose-response before being evaluated in an orthogonal chromatography-based GST-pulldown assay using cell lysates with over-expressed full length GST-MSN and Venus-Flag-CD44 (Fig. 4B). Hits confirmed in both assays were then assessed for binding to MSN by biophysical methods (see below). Out of 271 primary hits, two compounds with distinct chemical scaffolds (Fig. 4C) were selected to be evaluated further.

**Figure 4:**
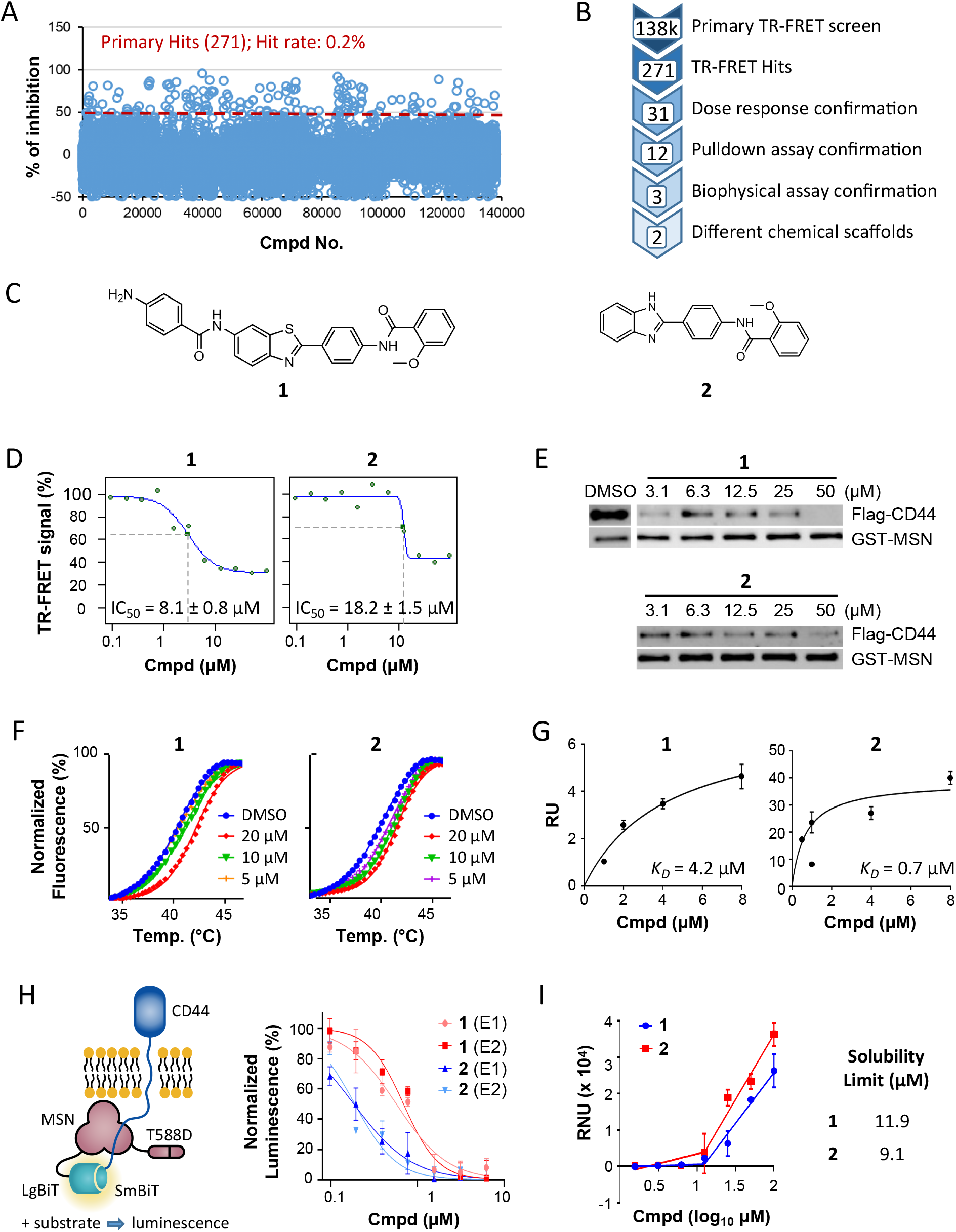
uHTS of a 138k compound library identifies MSN–CD44 small-molecule PPI inhibitors. (**A**) Scatter plot showing percentage inhibition with 138,214 compound library in the MSN–CD44 TR-FRET assay. Primary hit rate is 0.2%, with 271 compounds giving >50% inhibition. (**B**) Screening flow chart showing hit-to-lead identification strategy. (**C**) Structures of lead compounds based upon different chemical scaffolds. (**D**) Confirmatory TR-FRET dose response curves. (**E**) Dose response of compounds in GST pull down assay. Cell lysates were obtained from cells transfected with full length constructs of GST-tagged MSN and Flag-tagged CD44, and were incubated with the compounds. Immunoblotting was performed to detect GST and Flag. (**F**) Thermal shift assay, showing melting temperature (*T*_*m*_) curves of MSN with compounds. The average change in *T*_*m*_ (av. Δ*T*_*m*_), compared to DMSO alone, was determined from two experiments (**1**: 1.1 °C at 20 µM; **2**: 1.9 °C at 20 µM). (G) Dose response of compound binding to MSN FERM domain, measured by SPR. *K*_*D*_ measurements were determined for compounds **1** (4.2 µM) and **2** (0.7 µM). RU=response units (**H**) Dose response of compounds in cell-based NanoBiT bioluminescence assay, with HEK293 cells expressing LgBiT-MSN(T588D) and SmBit-CD44 fusions. Luminescence from reconstituted split luciferase was measured after substrate addition. Activities were determined from 2 independent experiments (**1**: 0.57±0.17 and 0.69±0.12 µM; **2**: 0.18±0.04 and 0.19±0.06 µM). (**I**) Solubility of compounds (PBS buffer, 1% DMSO), as measured by nephelometry. RNU=relative nephelometric units.

In the primary TR-FRET PPI assay, compounds **1** and **2** showed inhibitory values in the 5–20 μM range (Fig. 4D). Dose response analysis of the compounds in the GST-pulldown assay revealed that they were also active against full length proteins in cell lysates (Fig. 4E). Orthogonal biophysical assays were then performed on the compounds to confirm their activity. Both compounds could bind MSN in a thermal shift assay, stabilizing the protein by 1–2 °C (Fig. 4F). Moreover, surface plasmon resonance (SPR) experiments showed that binding of **1** and **2** to MSN could be fitted to a 1:1 model, with 4.2 and 0.7 μM *K*_*D*_ values, respectively (Fig. 4G).

To determine whether the identified compounds could interfere with MSN binding to CD44 in live cells, we established a split luciferase (NanoBiT) bioluminescence assay (Fig. 4H, left panel). MSN was fused to the large BiT at the N-terminus, while CD44 was fused to the small BiT at the C-terminus, with a constitutively active form of MSN (T588D) used to improve luminescence. Compounds **1** and **2** were found to be potent inhibitors of CD44–MSN interaction in live cells, with IC_50_ values of approximately 600 and 200 nM, respectively (Fig. 4H, right panel). Co-crystallization attempts of compound **1** and **2** with MSN were unsuccessful, likely because the compounds have low solubility in aqueous buffers (Fig. 4H).

### Structure-activity relationship studies with primary MSN–CD44 PPI hits

In order to improve the potency, selectivity, and solubility of compounds **1** and **2**, we conducted structure–activity relationship (SAR) studies using compounds that we synthesized in-house. We tested the derivatives in the MSN–CD44 TR-FRET assay, and also utilized a TR-FRET counter screen employing identical FRET pairs in an unrelated PPI (SYK-ITAM) to eliminate false hits that interfere with the assay or are protein aggregators. We synthesized 17 derivatives of compound **1** while retaining the 2-phenylbenzothiazole core (Table S2), but most of the benzothiazole derivatives either registered strongly in the TR-FRET counter-screen or exhibited no activity. Therefore, we decided to focus on compound **2** SAR. We synthesized twelve derivatives of compound **2**, all retaining the N-(4-(benzimidazole-2-yl)phenyl)acetamide core (Table S3). Of these, the 2,4-methoxybenzamide (**2a**) and the 3-chlorobenzamide (**2b**) substituents exhibited an 8-fold and 2-fold increase in potency, respectively, while displaying no activity in the counter screen (Fig. 5A). Furthermore, both compounds **2a** and **2b** showed improved solubility in PBS buffer when compared to compound **2**. Further testing demonstrated that compounds **2a** and **2b** were able to inhibit the MSN–CD44 interaction in cell lysates (Fig. 5B). Additionally, both compounds could stabilize MSN by 1–2°C in thermal shift assays at low micromolar concentrations (Fig. 5C).

**Figure 5:**
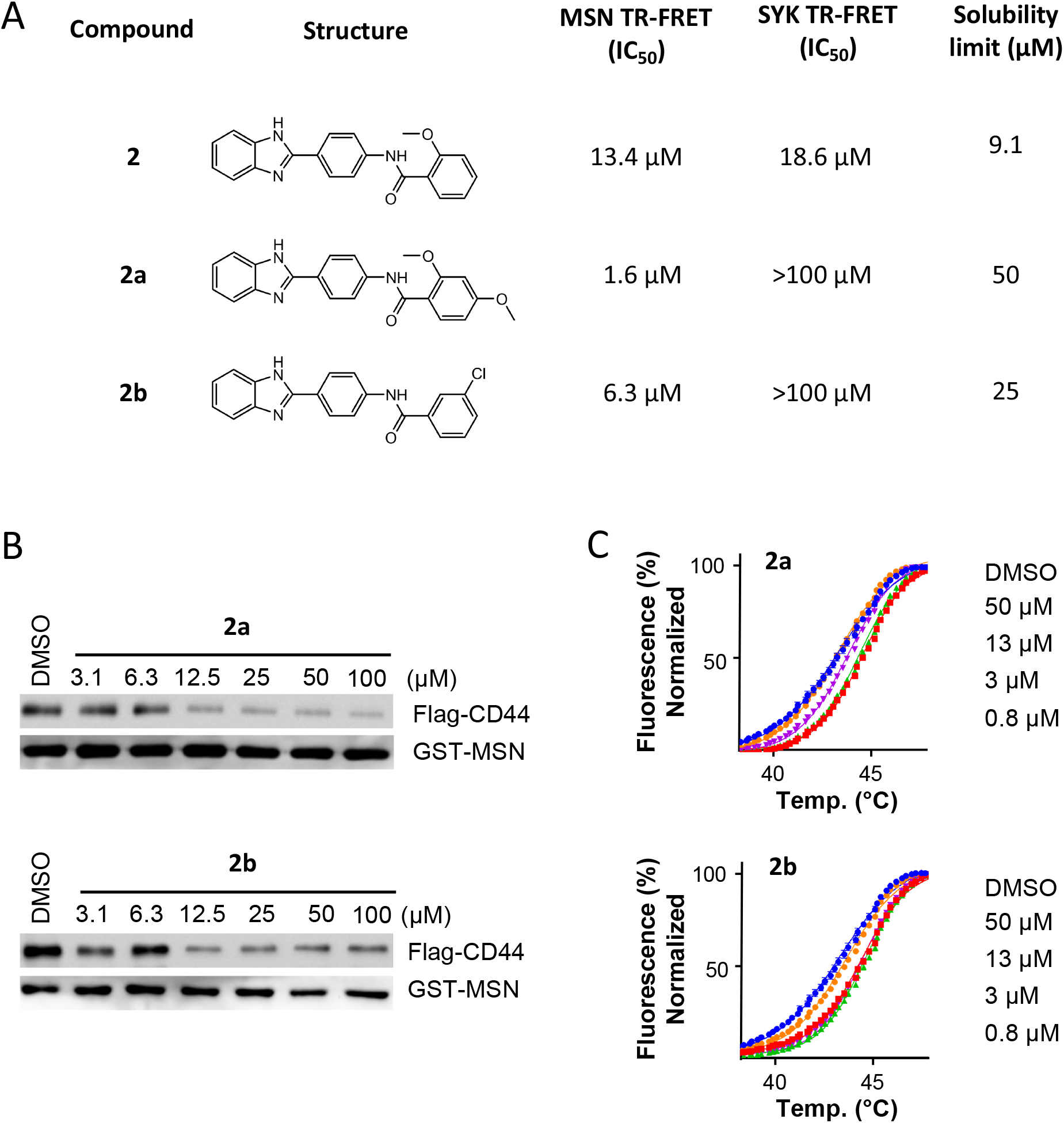
Compound **2** and analogues disrupt the MSN–CD44 PPI. (**A**) Structures of compound **2** and analogues, with associated solubility data (PBS, 1% DMSO), and activity in both the MSN–CD44 TR-FRET assay and an unrelated PPI (SYK-FCER1G) TR-FRET assay. Solubility was measured by nephelometry. (**B**) Dose response of compounds in GST pull down assay. Cell lysates were obtained from cells transfected with full length constructs of GST-tagged MSN and Flag-tagged CD44, and were incubated with the compounds. Immunoblotting was performed to detect GST and Flag, and a representative immunoblot from two experiments is shown. (C) Thermal shift assay, showing representative melting temperature (*T*_*m*_) curves of MSN with compounds. The average change in *T*_*m*_ (av. Δ*T*_*m*_), compared to DMSO alone, was determined from two experiments (**2a**: 1.3 °C at 50 µM; **2b**: 1.2 °C at 50 µM).

## Discussion

Proteomic data from postmortem frontal cortex samples in AD cases and controls have been used in network modelling to identify two proteins, MSN and CD44, which are co-regulated and linked to disease progression, including cognitive and pathological features (6). Although the impact of elevated levels of these proteins in AD brains is uncertain, inhibiting their function in microglia may reduce inflammation and prevent neuronal damage. Since CD44 is known to bind to the FERM domain of MSN, small molecule inhibitors that prevent this protein–protein interaction could potentially be useful tools for testing this hypothesis. Since MSN, along with other ERM proteins, is involved in critical cellular processes like cell adhesion, migration, and signalling (12), inhibiting a specific PPI with CD44 may be better tolerated by cells and impact only a portion of its functions. Although FERM domain proteins are known to be involved in various diseases, including cancer, cardiovascular, neurological, and immunological disorders (23-25), there is currently limited knowledge on the feasibility of developing small-molecule modulators that can target these proteins for therapeutic interventions. In one study, screening of the ERM protein ezrin against a compound library identified binders that inhibited formation of a phosphorylated active form of the protein (26), although the mechanism by which they blocked phosphorylation is not clear. In this work, we show that the FERM domain is a tractable target. Small molecule inhibitors with moderate potency were identified that could prevent the association of the FERM domain of MSN to the receptor tail of the CD44. Furthermore, by using phage display, peptide binders were found that could modify the association of MSN with CD44 via an allosteric mechanism, highlighting other locations for compound binding that impact the functionality of the FERM domain.

Structural studies on the MSN FERM domain (this work and (21)) show that ERM proteins can adopt either an “open” or “closed” conformation, the open form of which is permissive for binding of receptor tails, independent from the inhibitory C-terminal tail. Comparison of our peptide bound and ligand free MSN FERM domain structures show that the β5–β7 beta-sheet with the F3 lobe is displaced from the extended alpha helix by 2 Å in order to accommodate the C-terminal tail of CD44. Through binding to MSN, CD44 makes extensive contacts to the beta sheet, forming an antiparallel beta strand. Mutagenesis and structural data point to CD44 making hydrophobic contacts with most exposed hydrophobic residues within the binding pocket. In addition, CD44 makes contact with the extended alpha helix. In particular, interaction of H288 within the helix with the peptide backbone was critical for CD44 interaction. We propose that small molecules that bind either to the hydrophobic pocket within MSN or interact with H288 may interfere with CD44 binding.

During the phage display screening for allosteric pockets on the MSN FERM domain, we identified a peptide (C3P-pd) that binds to a cleft situated between the F1 and F3 subdomains, close to the suggested PIP_2_ binding site. The binding of the peptide caused the F3 lobe to assume an “open” conformation permissive to receptor binding. A previous study by Hamada et al. (2000) reported a similar “open” structure of radixin when it was co-crystallized with IP_3_, the head group of PIP_2_ (16). This led to the proposal that PIP_2_ binding to full-length ERM proteins helps to displace the inhibitory C-terminal domain through a shift of the beta-sheet in the F3 lobe. However, a later study found that PIP_2_ is required for the binding of MSN to the C-terminal tails of receptors, even when the C-terminal domain is disengaged through the use of the T558D phosphomimetic mutation (27). Thus, we suggest that PIP_2_ may induce the F3 subdomain to adopt an “open” state, which can enhance receptor binding in addition to or instead of helping to disengage the C-terminal domain. However, it is important to note that C3P-pd may not stimulate binding within the F3 subdomain in the same way as PIP_2_. C3P-pd did not enhance CD44 binding, and actually inhibited binding of a second phage display peptide (C3S1-pd) that bound similarly to CD44. Interestingly, C3P-pd binding was stimulated by C3S1-pd, suggesting that the allosteric modulation works in both directions. It is conceivable that receptor tail binding at the membrane may stimulate PIP_2_ binding as well. In summary, our findings demonstrate that ligand binding in the F1-F3 cleft can alter receptor tail binding. The discovery of C3P-pd could potentially lead to the development of peptidomimetics that allosterically regulate CD44 binding to ERM proteins. It would be valuable to investigate if C3P-pd can compete with PIP_2_ for MSN binding, and if this is the case, designing peptidomimetics based on C3P-pd could block PIP_2_ binding to ERM proteins.

Our study aimed to identify small molecule inhibitors of the MSN–CD44 interaction. High-throughput screening of a large chemical library revealed two hit series with distinct chemotypes. Both hits successfully displaced CD44 from MSN, with IC_50_ values in the low micromolar range, in both experiments with purified proteins and cell lysates. Moreover, the compounds exhibited potency in live cells. It remains unclear why these compounds show greater effectiveness in a cellular environment compared to when used with isolated proteins and peptides. One possible explanation is that the use of full-length protein in the context of membrane association enhances compound potency. Alternatively, the compounds may undergo metabolic activation within the cell. We cannot exclude off-target effects for the observed increase in activity in cells. We attempted to perform initial SAR on the thiazole hit but failed to obtain significant improvements in activity. Nevertheless, we made progress with the benzimidazole hits by enhancing potency while minimizing activity in a counter screen. Although compound solubility in aqueous buffers of this series remains a challenge, our SAR experiments indicated that increased solubility does not necessarily go hand-in-hand with decreased potency. We are currently evaluating the potential of these hits to alleviate microglial hyperactivity in tissue culture assays.

In summary, our findings provide a solid foundation for future medicinal chemistry efforts targeting ERM proteins for AD therapy. We have demonstrated that FERM proteins are a feasible target for the development of protein–protein interaction inhibitors. However, further optimization of the lead compounds is necessary to improve their potency and physiochemical properties as well as to evaluate their suitability for use in cellular assays.

## Experimental procedures

### DNA constructs

Expression plasmid constructs used in protein purification were generated by PCR amplification of the FERM domain of human MSN (residues 1-345) from the Mammalian Gene Collection cDNA library (IMAGE:4908580). The PCR product was cloned into the *E. coli* expression vector pNIC28-Bsa4 using ligation independent cloning, giving rise to an N-terminal fusion with a 6His tag and a TEV protease cleavage site (MSNA-c001). Constructs for purification of MSN-H288A (MSNA-c028), MSN-L281R (MSNA-c025), and the MSN-H288A, -L281R (MSNA-c033) were produced by site-directed mutagenesis, using the KLD enzyme kit (NEB, M0554S). Full-length Venus-Flag-tagged CD44 and GST-tagged MSN for mammalian expression were generated using Gateway© cloning kit (Invitrogen). The vector backbones are pDEST27 vector (Invitrogen) for GST-tagged MSN and lab customized pSCM167 for Venus-Flag-tagged CD44. For NanoBiT split luciferase assay, BiBiT Flexi vectors were custom ordered from Promega.

### Protein purification

Wild-type and mutant MSN were produced using *E. coli* BL21(DE3)-R3-pRARE cells grown in terrific broth. Prior to harvesting, cells were shifted to 18 °C for 16 h after IPTG induction. Cells were lysed by sonication in Lysis Buffer containing 50 mM HEPES (pH 7.5), 500 mM NaCl, 10 mM imidazole, 5% glycerol, 1 mM TCEP. MSN protein was bound to equilibrated Ni-IDA resin for 1 h before two batch washes in Lysis Buffer. Beads were subsequently loaded onto a drip column, followed by washing with Lysis Buffer containing 30 mM imidazole, before finally eluting bound protein with Lysis Buffer containing 300 mM imidazole. For protein crystallography, the 6His tag was cleaved off by the addition of a 1:10 mass ratio of 6His-TEV protease while undergoing dialysis (10k MWCO) in Lysis Buffer lacking imidazole for 16h at 4 °C. TEV protease and the cleaved 6His tag was subsequently removed with Ni-IDA resin equilibrated in Lysis Buffer. For TR-FRET assays, the N-terminal tag was left intact. Both tagged and untagged forms of MSN were further purified by SEC in Lysis Buffer lacking imidazole. The molecular mass of purified proteins was confirmed by intact mass spectrometry.

### Crystallization and data collection

Crystals of the FERM domain of either wild-type or H288A mutant MSN (1-345) were obtained by first incubating protein with a 21-residue CD44 peptide (672-SRRRCGQKKKLVINSGNGAVEDY-693) at a 1:1.1 ratio (final 12-14 mg/ml protein concentration) for 30 min on ice. Wild-type and H288A mutant proteins were crystallized at 4 °C in sitting drops by addition of reservoir (0.1-0.2M ammonium acetate, 32-34% 2-propanol, 0.1M Tris pH8.5) in 1:2 or 2:1 ratios, respectively. Although the CD44 peptide was necessary for crystallization of these proteins, the peptide was not visible in structures obtained. The MSN-L281R mutant was crystallized in sitting drops containing a 1:2 ratio of protein to reservoir (20% PEG3350, 0.2M NaF, 10% ethylene glycol, 0.1M bis-tris-propane pH 8.5) at 4 °C. Crystals of MSN (1-345) with bound CD44 were obtained by first incubating MSN with a minimal 8-residue CD44 peptide (678-QKKKLVIN-685) at a 1:5 ratio (final 16 mg/ml protein) for 30 min on ice, followed by incubation in drops containing a 1:1 ratio of protein-peptide mix to reservoir (20% PEG3350, 0.2M potassium thiocyanate, 10% ethylene glycol, 0.1M bis-tris-propane pH 6.5) at 4 °C. Crystals of MSN (1-345) with bound phage display peptide C3P-pd were obtained by incubating MSN with peptide (EDGGSWKYPDAFELSG) at a 1:2 ratio (final 18 mg/ml protein) for 30 min on ice, followed by incubating drops containing a 1:2 ratio of protein-peptide mix to reservoir (20% PEG3350 10% ethylene glycol, 0.2M NaBr, 0.1M bis-tris-propane pH 7.5) at 4 °C. Crystals of MSN (1-345) bound to both phage display peptides C3P-pd and C3S1-pd (EDGGSSWEYIWTLPSG) were obtained by incubating protein and peptides in a 1:2:2 ratio (final 16 mg/ml protein) for 30 min on ice, followed by incubating drops containing a 2:1 ratio of protein-peptide mix to reservoir (25% PEG4000, 0.2M ammonium sulfate, 0.1M acetate pH 4.5) at 4 °C. All crystals were grown in 150 nL sitting drops. Most crystals were cryoprotected by the addition of 1 μL of reservoir solution immediately before harvesting. For crystals grown in 2-propanol, 1 μL of 25% PEG 3350 was used.

Data were collected at Diamond Light Source beamlines I03 and I04 and indexed integrated with Dials (28). Data with low anisotropy (6TXQ, 6TXS, 8CIR) were scaled with Aimless (29), while data with significant anisotropy were scaled with Staraniso (30) and Aimless. The anisotropic high resolution cut-offs were as follows: 8CIS: 2.12-1.42 Å, 8CIT: 3.97-2.54 Å, 8CIU: 3.24-2.39 Å. The Apo structure was determined first by molecular replacement performed by Phaser (31) using 1E5W as a model (21). The apo structure was used in molecular replacement to determine the remaining structures. The two mutant structures were refined with successive rounds of Coot (32) and Buster (33), while the other structures were refined with Coot and Refmac5 (34). Waters were initially modelled in the L281R structure (8CIT) and did cause a decrease in R-factors, but the majority refined to abnormally low B-factors that are assumed to have been a result of noisy, anisotropic data with low completeness at a resolution required to see waters. It was decided that all waters would be removed for the deposition. The final models were verified with MolProbity (35) and deposited in the PDB under the accession codes shown in Table S1. Movement of FERM subdomains was analysed by DynDom (36).

### TR-FRET assay

MSN TR-FRET experiments were performed in 20 μl reactions, using 384-well shallow-well microplates. Final reaction components contained 6His-MSN (2 nM), FITC-conjugated CD44 peptide (8 nM; SRRRCGQKKKLVINSGNGAVEDYK-FITC), Tb-conjugated anti-6His antibody (53 ng/ml; Cisbio) in assay buffer containing 25 mM HEPES (pH 7.5), 200 mM NaCl, 0.1% BSA and 0.05% Tween-20. After a 2h incubation (at RT), TR-FRET signals were measured on a BMG Labtech PHERAstar FSX reader using a Lanthascreen Optics Module. A 200 µs delay was used after excitation with a flash lamp before measurement of fluorescence emission at 490 and 520 nM. TR-FRET ratios of fluorescent intensity at 520 nm to 490 nm were calculated. The half maximal inhibitory concentration (IC_50_) was determined by fitting a four parametric logistic curve to the data.

### Phage display screening

MSN was screened against an unbiased, random 10-mer peptide phage display library utilizing the pIII coat protein of filamentous M13 phage, as previously described with modification (37, 38). Briefly, His-bind magnetic Dynabeads (Invitrogen) were washed with TBS containing 0.05% Tween-20 (TBS-T) and then incubated with either 50 µl of 3.2 µM 6His-MSN in TBS-T or buffer alone for 30 min at RT with rotation. All selections were carried out in low-retention microfuge tubes. The beads were washed and blocked 3 times with casein blocking buffer (CBB; 1% Hammarstein-grade casein in Tris-buffered saline, pH 7.4), and MSN-beads were incubated for 30 min at RT with or without of 1 µM CD44 competitor peptide. The phage library, diluted in CBB, was added to give 5 × 10^11^ phage molecules. Selections were incubated for 2h at 4 °C with rotation followed by four washes with TBS-T. Bound phage were eluted with 100 µl of 0.2M glycine (pH 2.5) for 8 min at 37 °C, followed by immediate neutralization with the addition of 10 µl of 1 M Tris (pH 11). Phage selections were amplified by adding them to 1.0 ml TG1 cells (Lucigen) grown to mid-log phase (OD ∼ 0.6) and incubated at 37 °C for 1.5 h. Subsequently, cells were diluted 1:50 into 30 ml 2xYT media containing 15 µg/ml tetracycline and grown overnight at 37 °C. At the same time, phage were titered using TG1 cells and tetracycline agar plates. The following day, phage were precipitated with 5X PEG 8000-NaCl (20% PEG/2.5 M NaCl) and pelleted phage were washed with and resuspended into TBS. Selections were repeated three times with input phage dropped 5-fold for each round, resulting in enrichment (>10,000-fold) over control selections with only beads. Phage isolated from individual colonies were screened for MSN binding using ELISA. Briefly, 0.5 µg MSN in PBS was adsorbed to a 384-well white Nunc® MaxiSorp™ plate overnight at 4 °C. Phage were serially diluted 3-fold in CBB and added to coated wells. The plate was incubated for 1.5h at RT, and unbound phage were removed by washing 3 times with TBS-T. A 1:10,000 dilution of anti-M13-HRP in CBB was added to each well and incubated for 1h at RT. Wells were washed 3 times with TBS-T and phage binding was detected by addition of 20 µl SuperSignal™ ELISA Pico Chemiluminescent Substrate (Thermo-Fisher Scientific) and read on an Envision multilabel plate reader (Perkin-Elmer). Phage ELISA binding curves were analysed using a four-parameter non-linear curve fit to determine phage EC_50_.

### Ultra-high-throughput screening

Ultra-high throughput screening (uHTS) for small molecule PPI inhibitors of MSN–CD44 interaction was performed using the TR-FRET assay miniaturized into the 1536-well format. Briefly, the optimized reaction mixture (1.5 nM of 6His-MSN and 25 nM of FITC-CD44 peptide in the presence of anti-His-Tb at final 1:1000 dilution) was dispensed at 5 µL per well into 1536-well black plates (Corning Costar, #3724). 0.1 µL of library compound dissolved in DMSO was added using a pin tool integrated with the Biomek NX automated liquid handling workstation (Beckman Coulter). Final compound and DMSO concentrations were 20 µM and 2%, respectively. The plates were centrifuged at 1000 rpm for 5 min and incubated at RT for 2 h. TR-FRET signals were then measured using the BMG Labtech PHERAstar FSX reader with the HTRF optic module (excitation at 337 nm; emission A at 490 nm; emission B at 520 nm). TR-FRET ratios of fluorescent intensity at 520 nm to 490 nm were calculated.

Screening data were analyzed using Bioassay software from CambridgeSoft (Cambridge, MA). The singal-to-background (S/B) ratio and Z’ for each screening plate were calculated using the following equations: S/B = F_control_/F_blank_, Z’ = 1-[(3*SD_control_ + 3*SD_blank_)/ (F_control_ - F_blank_)]. The inhibitory effect of compounds on the TR-FRET signal was expressed as a percentage of control, and calculated on a plate-by-plate basis, as follows: % of Inhibition = 100 – [(F_compound_ - F_blank_)/(F_control_ - F_blank_)*100], where F_compound_ is the TR-FRET signal from compound wells, F_control_ is the average TR-FRET signal from DMSO control wells, which defines the highest signal, and F_blank_ is the average signal from MSN and anti-His-Tb only wells, lacking the FITC peptide. Compounds where percentage inhibition was greater than 50 were defined as primary hits. The S/B across all screening plates was greater than 10, and the Z’ greater than 0.5. A chemical diversity library containing 138,214 compounds (ChemDiv and Asinex) was screened. After cherry-picking the hits from the library stock and confirming their activity through dose-response in the TR-FRET assay, the primary hits were repurchased for further confirmation by dose-response in the TR-FRET assay.

### GST-pull down assay

TR-FRET-confirmed hits were validated in a GST-pull down assay using cell lysates of HEK293 cells co-expressing full-length constructs of both Venus-Flag-tagged CD44 and GST-tagged MSN. The Venus tag was used as a marker for CD44 protein expression. Dose responses were performed by the addition of 0.5 μl of compounds (in DMSO) to 100 μl of cell lysates which were diluted in 0.25% triton lysis buffer (150 mM NaCl, 10 mM HEPES, pH 7.5, 0.25% Triton X-100, phosphatase inhibitor (Sigma, P5726) and protease inhibitor cocktail (Sigma, P8340)). The mixtures were rotated for 1 h at 4 °C before the addition of 15 μL of 50% pre-equilibrated GST resin. After incubation on a rotator for 1.5 h at 4 °C, the GST resin was washed three times with 1 ml of 0.25% triton lysis buffer. SDS-sample buffer was added prior to boiling for 5 min. Samples were analysed by SDS-PAGE and immunoblotting, using anti-FLAG antibody for CD44 (1:1000 dilution of anti-Flag M2-HRP antibody; Sigma, A8592) and anti-GST antibody for GST-MSN (1:1000 dilution of anti-GST (2.6H1) Mouse mAb; Cell Signaling, 2624S). For GST detection, a secondary HRP-conjugated antibody was used (1:5000 dilution of AffiniPure Goat Anti-Mouse IgG (H+L); Jackson Immuno Research, 115-035-003).

### Thermal shift assay

Compounds in DMSO were diluted 1 in 50 in assay buffer (25 mM Tris pH 8.0, 150 mM NaCl, 5% glycerol) containing 2 µM MSN. After removal of compound precipitates by centrifugation, the mixture was incubated for 30 min at RT. Sypro Orange (Thermofisher S6650) was then added at 20x concentration. Melting curves were obtained on a qPCR machine (Eppendorf RealPlex 4), ramping up from 25 to 95 °C, at 1 °C min^−1^. To calculate the T_m_, data was fitted to the Boltzmann equation using Prism GraphPad.

### Surface plasmon resonance assay (SPR)

Binding experiments were performed using a Biacore X100 instrument with a carboxymethylated dextran (CM5) sensor chip. Protein ligand was bound to the sensor surface by amine coupling in acetate buffer pH 4, giving 4000 response units (RU). The protein-bound sensor was equilibrated in running buffer (20 mM HEPES pH 7.5, 200 mM NaCl, 1 mM TCEP, 0.005% tween-20) prior to compound injection. Multi-cycle kinetic analysis was performed with compounds (0.5-8 μM). Data were fitted with the Biacore evaluation software, using a 1:1 binding model.

### Isothermal titration calorimetry (ITC)

ITC was performed to quantify the thermodynamics of binding between MSN and the synthetic phage display peptides. MSN was buffer exchanged (NAP-5 column; GE Healthcare Life Sciences) and diluted to 20 µM in ITC buffer (50 mM HEPES, pH 7.4, 200 mM NaCl, and 0.5 mM TCEP). Phage display peptides C3P-pd and C3S1-pd were diluted to 200 µM in ITC buffer. Protein and peptide concentrations were calculated by measuring their absorption at 280 nm and using their expected extinction coefficients, calculated by Protparam (https://web.expasy.org/protparam/). ITC measurements were recorded at 25 °C using an AutoITC 200 microcalorimeter (Malvern Instruments). Injections of 0.2 µl peptide were titrated into 200 µl MSN at 180s intervals with a reference power of 7 µcal/sec. Estimated heats of dilution were subtracted and titration data analyzed using Microcal LLD Origin7 software by non-linear least squares, and fitting a one-site model as a function of the peptide:protein ratio.

### NanoBit split luciferase assay in live cells

The HEK293T (ATCC, CRL-3216) cells were grown in Dulbecco’s modified Eagle’s medium (DMEM; Corning, 10-013-CV) supplemented with 10% fetal bovine serum (Sigma, F0926-500ML), 100 U/ml penicillin, and 100 µg/ml streptomycin (Sigma, P0781-100ML) at 5% CO2 and 37°C in a humid environment. Four BiBit Flexi vectors for each of the full-length CD44, MSN(T558D), and MSN(T558A) genes with either N-terminal or C-terminal SmBit and LgBiT fusions were produced using Flexi®-based cloning (Promega). Best luminescence signals were obtained with CD44 containing a C-terminal smBiT fusion (C-terminal smBiT) and MSN(T558D) containing an N-terminal LgBiT fusion (C-terminal smBiT). LgBiT-MSN(T558A) was used as negative control. To evaluate the effects of the compounds, 30 μL of HEK293T cell suspension was seeded at 6000 cells/well in a 384-well plate (Corning, 3826). The next day, a mixture of 20 ng of CD44-smBiT and 5 ng of LgBiT-MSN(T558D) plasmids was added to 75 ng of Fugene HD (Promega, Cat#2312) transfection reagent diluted in DMEM, and 10 μl of the mixture was added to each well. After 18 hours, the cells were treated with a 2-fold serial dilution of compounds (6.25-0.098 μM) that was dissolved in DMSO and then incubated for 3 hours. Next, 10 μl of Nano-Glo® Live Cell Reagent (Promega, N2014) was added to the wells, and the plates were incubated for 20 minutes at 37°C. The signal was then measured using the 2102 Envision Multilabel Reader (PerkinElmer).

### Synthesis details for selected compounds

Synthesis of compounds **1** (Scheme 1) and **2**, including analogues **2a** and **2b** (Scheme 2), are detailed below. Compound identity and purity were confirmed by NMR and LCMS, shown in supplementary information.

**Scheme 1.**
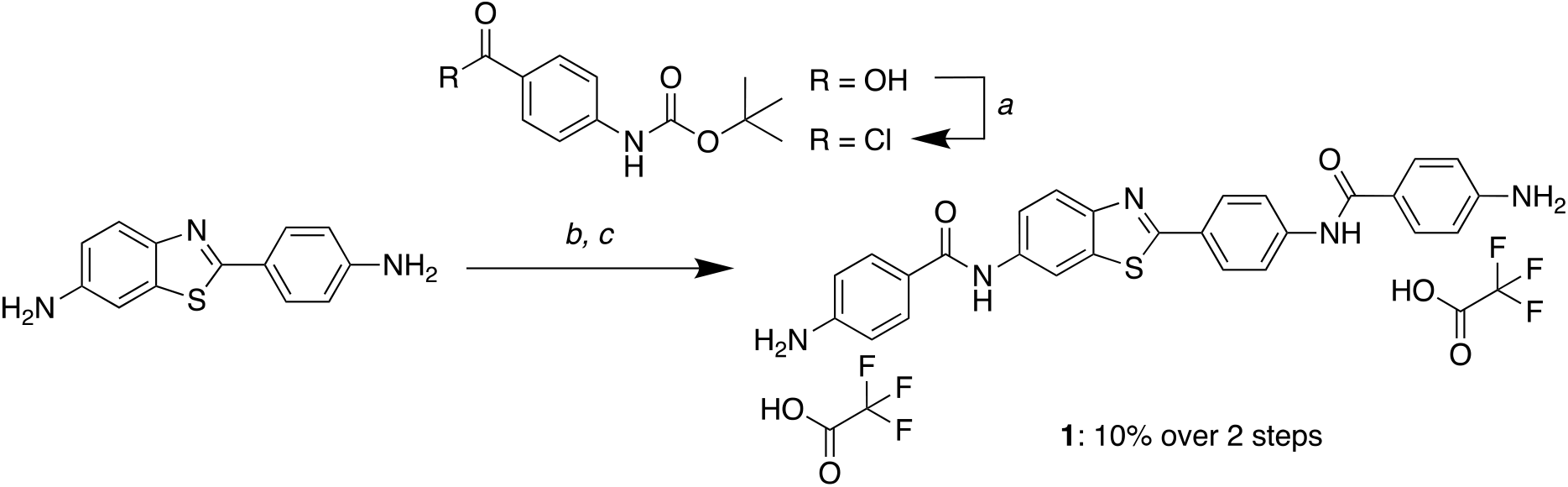
Synthesis of compound **1**. (a) 4-((*tert*-butoxycarbonyl)amino)benzoic acid, CH_2_Cl_2_, SOCl_2_, 0–40 °C; (b) *tert*-butyl (4-(chlorocarbonyl)phenyl)carbamate, DIPEA, THF, 65 °C; (c) TFA, CH_2_Cl_2_.

**Scheme 2.**
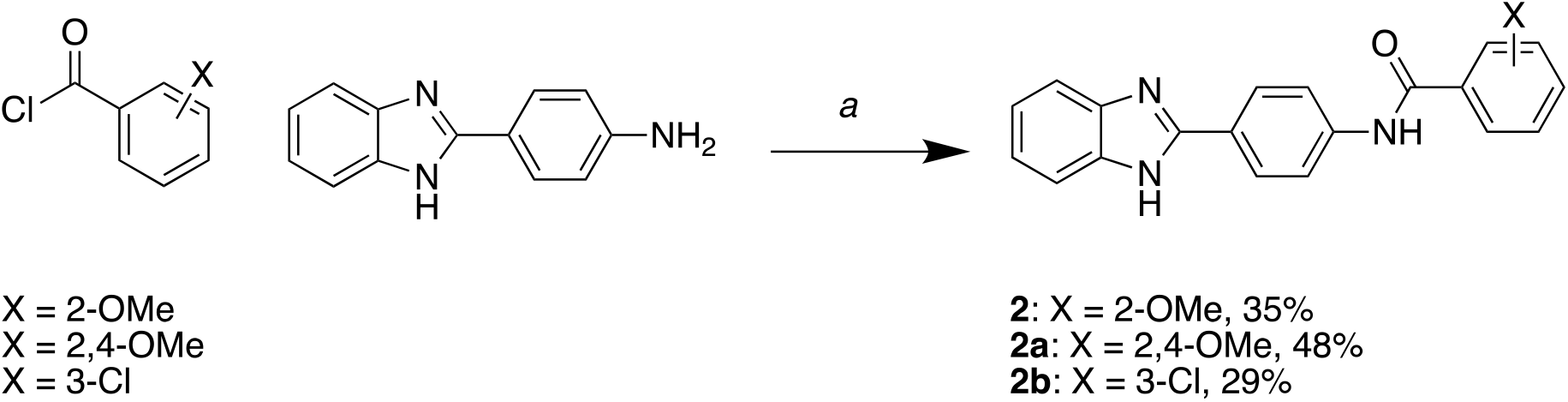
Synthesis of compounds **2, 2a** and **2b**. (a) Acyl chloride, DIPEA, THF, 0–65 °C.

#### Preparation of 4-amino-*N*-(4-(6-(4-aminobenzamido)benzo[*d*]thiazol-2-yl)phenyl)benzamide bis(2,2,2-trifluoroacetate) (1)

To a flask was added 4-((tert-butoxycarbonyl)amino)benzoic acid (236 mg, 2.4 equiv, 995 µmol) and CH_2_Cl_2_ followed by addition of thionyl chloride (103 µL, 3.4 equiv, 1.40 mmol) at 0 °C. The reaction was allowed to warm to rt and then stirred at 40 °C for 4 h. The reaction was concentrated *in vacuo* to yield a yellow solid. To this was added 2-(4-aminophenyl)benzo[*d*]thiazol-6-amine (100 mg, 1.0 equiv, 414 µmol) and DIPEA (217 µL, 3.0 equiv, 1.24 mmol) in THF (1.66 mL, 0.25 M) and the reaction stirred at 65 °C. After 1 h a precipitate crashed out of solution. The reaction was cooled and diluted with H_2_O and the precipitate was collected via vacuum filtration. The product was further purified by washing with CH_2_Cl_2_ under vacuum to yield the desired product *tert*-butyl (4-((2-(4-(4-((*tert*-butoxycarbonyl)amino)benzamido)phenyl)benzo[*d*]thiazol-6-yl)carbamoyl)phenyl)carbamate as a white solid, which was used without further purification. *Tert*-butyl (4-((2-(4-(4-((*tert*-butoxycarbonyl)amino)benzamido)phenyl)benzo[*d*]thiazol-6-yl)carbamoyl)phenyl)carbamate was added to a vial with CH_2_Cl_2_ and TFA (1:1) and the reaction stirred at rt for 1 h. The reaction was concentrated *in vacuo* to yield the desired product 4-amino-*N*-(4-(6-(4-aminobenzamido)benzo[*d*]thiazol-2-yl)phenyl)benzamide bis(2,2,2-trifluoroacetate) as a yellow amorphous solid (28 mg, 10% over 2 steps). ^1^H NMR (400 MHz, DMSO-*d*_*6*_): δ 10.07 (d, *J* = 12.1 Hz, 2H), 8.62 (d, *J* = 2.0 Hz, 1H), 8.06 – 7.92 (m, 5H), 7.78 (dd, *J* = 8.7, 1.8 Hz, 5H), 6.71 – 6.65 (m, 4H). HPLC purity: >95%; LCMS: calculated for C_27_H_22_N_5_O_2_S [M + H]^+^: 480.15. Found: 480.09.

#### Preparation of *N*-(4-(1H-benzo[*d*]imidazol-2-yl)phenyl)-2-methoxybenzamide (2)

A solution of 4-(1*H*-benzo[*d*]imidazol-2-yl)aniline hydrochloride (0.200 g, 1.0 equiv, 0.810 mmol) in THF (3.0 mL) at 0 °C was treated with DIPEA (0.280 mL, 2.0 equiv, 1.60 mmol), followed by dropwise addition of 2-methoxybenzoyl chloride (0.150 mL, 1.2 equiv, 0.980 mmol) over 5 mins. After 1 h at 0 °C, the reaction mixture was allowed to warm to rt, and then was heated to 65 °C with stirring for 12 h. The product crashed out of solution and was collected via vacuum filtration using ethanol, followed by diethyl ether to collect the desired product *N*-(4-(1*H*-benzo[*d*]imidazol-2-yl)phenyl)-2-methoxybenzamide as an off-white amorphous solid (100 mg, 35%). ^1^H NMR (400 MHz, DMSO-*d*_*6*_): δ 10.61 (s, 1H), 8.37 (d, *J* = 9.0 Hz, 2H), 8.07 (d, *J* = 8.9 Hz, 2H), 7.83 (dd, *J* = 6.1, 3.2 Hz, 2H), 7.66 (dd, *J* = 7.6, 1.8 Hz, 1H), 7.59 – 7.49 (m, 3H), 7.22 (dd, *J* = 8.5, 0.9 Hz, 1H), 7.10 (td, *J* = 7.5, 1.0 Hz, 1H), 3.92 (s, 3H). HPLC purity: >95%; LCMS: calculated for C_21_H_18_N_3_O_2_ [M + H]^+^: 344.14. Found: 344.04.

#### Preparation of *N*-(4-(1H-benzo[*d*]imidazol-2-yl)phenyl)-2,4-dimethoxybenzamide (2a)

A solution of 4-(1*H*-benzo[*d*]imidazol-2-yl)aniline hydrochloride (0.200 g, 1.0 equiv, 0.810 mmol) in THF (3.0 mL) at 0 °C was treated with DIPEA (0.280 mL, 2.0 equiv, 1.60 mmol), followed by dropwise addition of 2,4-dimethoxybenzoyl chloride (0.200 g, 1.2 equiv, 0.980 mmol) over 5 mins. After 1 h at 0 °C, the reaction mixture was allowed to warm to rt, and then was heated to 65 °C for 12 h. The product crashed out of solution and was collected via vacuum filtration with ethanol, followed by diethyl ether to yield the desired product *N*-(4-(1*H*-benzo[*d*]imidazol-2-yl)phenyl)-2,4-dimethoxybenzamide as an off-white amorphous solid (150 mg, 48%). ^1^H NMR (400 MHz, DMSO-*d*_*6*_): δ 10.33 (s, 1H), 8.25 (d, *J* = 8.7 Hz, 2H), 8.07 (d, *J* = 8.7 Hz, 2H), 7.83 – 7.74 (m, 3H), 7.51 (dd, *J* = 6.3, 3.1 Hz, 2H), 6.74 (d, *J* = 2.3 Hz, 1H), 6.70 (dd, *J* = 8.7, 2.3 Hz, 1H), 3.98 (s, 3H), 3.86 (s, 3H). HPLC purity: >95%; LCMS: calculated for C_22_H_20_N_3_O_3_ [M + H]^+^: 374.15. Found: 374.37.

#### Preparation of *N*-(4-(1H-benzo[*d*]imidazol-2-yl)phenyl)-3-chlorobenzamide (2b)

A solution of 4-(1*H*-benzo[d]imidazol-2-yl)aniline hydrochloride (0.200 g, 1.0 equiv, 0.810 mmol) in THF (3.0 mL) at 0 °C was treated with DIPEA (0.280 mL, 2.0 equiv, 1.60 mmol), followed by dropwise addition of 3-chlorobenzoyl chloride (0.130 mL, 1.2 equiv, 0.980 mmol) over 5 mins. After 1 h at 0 °C, the reaction mixture was allowed to warm to rt, and further to 65 °C with stirring for 12 h. The mixture was allowed to cool to RT and the solvent was removed *in vacuo*. The crude material was sonicated with water (2.0 mL), followed by addition of CH_2_Cl_2_ (4.0 mL) for 5 mins to give a solid. The solid was collected via vacuum filtration with diethyl ether to yield the desired product *N*-(4-(1*H*-benzo[*d*]imidazol-2-yl)phenyl)-3-chlorobenzamide as an off-white amorphous solid (83 mg, 29%). ^1^H NMR (400 MHz, DMSO-*d*_*6*_): δ 10.86 (s, 1H), 8.36 (d, *J* = 8.8 Hz, 2H), 8.13 (d, *J* = 8.8 Hz, 2H), 8.07 (t, *J* = 1.9 Hz, 1H), 7.98 (dt, *J* = 7.8, 1.4 Hz, 1H), 7.82 (dd, *J* = 6.1, 3.1 Hz, 2H), 7.71 (ddd, *J* = 8.0, 2.2, 1.1 Hz, 1H), 7.61 (t, *J* = 7.9 Hz, 1H), 7.53 (dd, *J* = 6.1, 3.1 Hz, 2H). HPLC purity: >95%; LCMS: calculated for C_20_H_15_ClN_3_O [M + H]^+^: 348.09. Found: 347.99.

## Supporting information

This article contains supporting information.

## Acknowledgements

We would like to thank all members of the Emory-Sage-SGC TREAT-AD center, the TREAT-AD advisory board members, and the NIA Scientific Officers, Lorenzo Refolo and Suzana Petanceska, for their advice and guidance on the study. We would also like to acknowledge Diamond Light Source for granting us beamtime (proposals mx19301 and mx28172), and the personnel at beamlines I03 and I04 for helping with crystal testing and data collection. We thank the CMD Biotechnology group for help with plasmid cloning, test expression, and mass spectrometry.

## Funding and additional information

The Target Enablement to Accelerate Therapy Development for Alzheimer’s Disease (TREAT-AD) Consortium was established by the National Institute on Aging (NIA). The research reported in this manuscript was led by the Emory-Sage-SGC TREAT-AD center and supported by grant U54AG065187 from the NIA. The content is solely the responsibility of the authors and does not necessarily represent the official views of the NIA. The Structural Genomics Consortium is a registered charity (number 1097737) that receives funds from AbbVie, Bayer Pharma AG, Boehringer Ingelheim, Canada Foundation for Innovation, Eshelman Institute for Innovation, Genome Canada, Genentech, Innovative Medicines Initiative (EU/EFPIA), Janssen, Merck KGaA Darmstadt Germany, MSD, Novartis Pharma AG, Ontario Ministry of Economic Development and Innovation, Pfizer, São Paulo Research Foundation‐FAPESP, Takeda, and Wellcome. The funders played no role in study design, data collection and analysis, decision to publish, or manuscript preparation.

## Conflict of Interests

The authors declare that they have no conflicts of interest with the contents of this article.

## Author contributions

Y. D., A. I., S. F., O. G., A. L., A. A., K. P., H. F., and V. K. contributed to the design and conception of the study. Y. D., W. B., T. L., J. A.-G., K. Q., F. B., A. S., F. N., and V. K. performed experiments, analyzed and interpreted data. V. K. drafted the manuscript, and all authors provided critical feedback and revisions. All authors approved the final version of the manuscript.

## Abbreviations

2xTY: 2x yeast extract tryptone
5xFAD: five familial alzheimer’s disease mutation
AD: Alzheimer’s disease
BiT: binary technology
CBB: casein blocking buffer
DMSO: dimethylsulfoxide
ERM: ezrin radixin moesin
FERM: 4.1B ezrin radixin moesin
GST: glutathione S-transferase
HTS: high throughput screening
IP_3_: inositol triphosphate
ITAM: immunoreceptor tyrosine-based activation motif
IDA: iminodiacetic acid
ITC: isothermal titration calorimetry
LgBiT: large binary technology
MWCO: molecular weight cut-off
PDB: the Protein Data Bank
pd: phage display
PIP_2_: Phosphatidylinositol 4,5-bisphosphate
PPI: Protein-Protein Interaction
RU: Response Units
RMSD: root-mean-square deviation
RNU: relative nephelometric units
SAR: structure-activity relationship
SEC: size exclusion chromatography
S/B: signal-to-background ratio
SmBiT: small binary technology
TBS: tris-buffered saline
TCEP: tris(2-carboxyethyl)phosphine
*T*_*m*_: melting temperature
TR-FRET: time-resolved fluorescence energy transfer
uHTS: ultra-high throughput Screening

## Figure Legends

**Figure S1**: Comparison of new and previously reported MSN structures reveals flexibility of FERM subdomain orientations and alternate F3 subdomain conformations.

(A) Overlay of our novel human MSN FERM structure (6TXQ) with a previous structure (1E5W). FERM subdomains of 6TXQ are coloured as in Fig. 1B, while 1E5W is coloured pink. (B) Top view of overlaid 6TXQ and 1E5W structures, with alignment over the F1 subdomain. The F3 subdomain in 6TXQ is rotated 12° along the axis of the long alpha helix, as measured by DynDom. (C) Overlay of individual subdomains from 6TXQ and 1E5W. Both F1 and F2 subdomains have small RMSD values and are closely aligned. In contrast, the F3 subdomain has a large RMSD value, with “open” and “closed” conformations observed for the previous (1E5W) and new (6TXQ) structures, respectively. RMSD=root mean square deviation.

**Figure S2**: Structures of MSN FERM domain containing either H288A or L281R point mutations show minimal differences to wild-type FERM domains.

The crystal structures of mutant FERM domains of human MSN, containing the point mutations H288A (PDB: 8CIU) or L281R (PDB: 8CIT). Shown are cartoon models, with the FERM subdomains coloured the same way as in Fig. 1B. The mutated residues are highlighted in magenta. The RMSD values of mutant FERM domains, compared to wild-type (PDB: 6TXQ), are low, showing minimal differences in protein folding. In the case of MSN-L281R, the RMSD value is an average from three molecules in the asymmetric unit. RMSD=root mean square deviation.

**Figure S3**: Binding kinetics of phage display peptides by ITC

(**A, B**) Example ITC isotherms of MSN FERM domain with peptide injections (upper) and integrated peak fit (lower) using a 1:1 binding model. (**A**) C3P-pd peptide. (**B**) C3S1-pd peptide.

## Notes

### Competing Interest Statement

The authors have declared no competing interest.

