## Supplemental Information for "Development of FERM domain protein-protein interaction inhibitors for MSN and CD44 as a potential therapeutic strategy for Alzheimer’s disease"

### Contents:

Figures S1-S3
Tables S1-S3
List of Emory-Sage-SGC TREAT-AD Center members
Chemical Synthesis: $^1$H NMR and LCMS analysis

### Figure S1

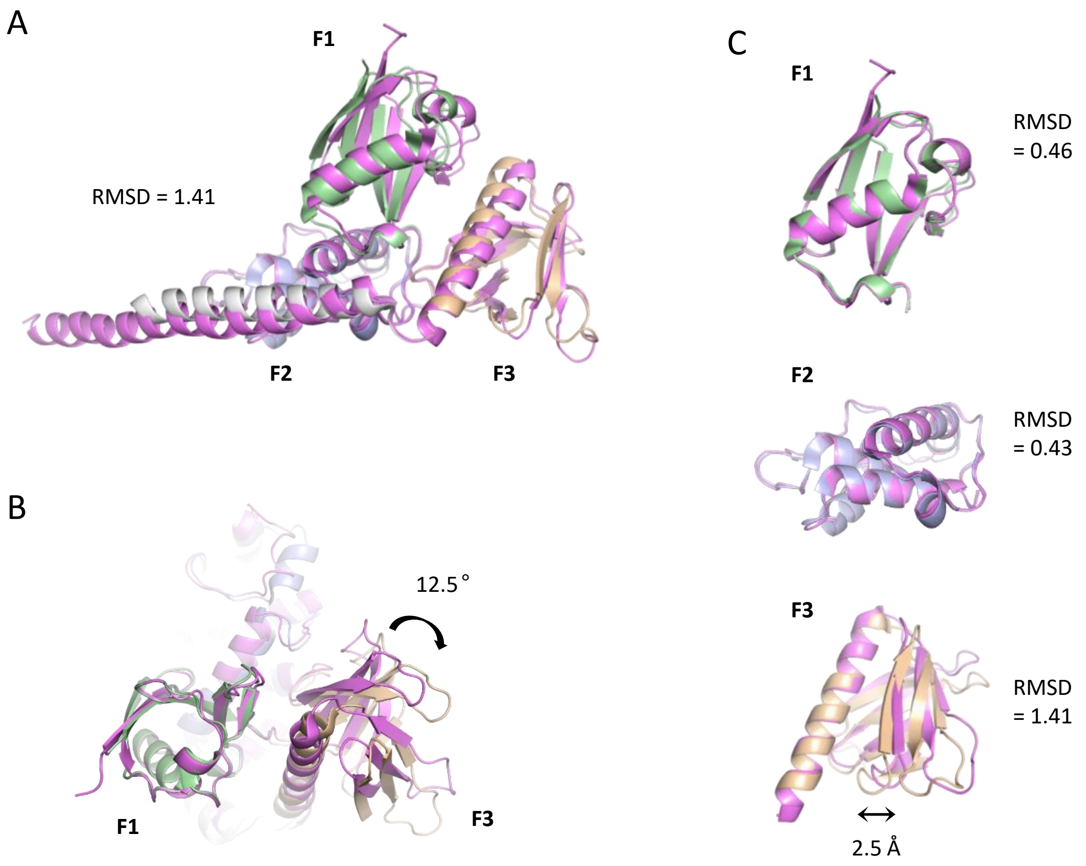


**Figure S1**: Comparison of new and previously reported MSN structures reveals flexibility of FERM subdomain orientations and alternate F3 subdomain conformations.

(A) Overlay of our novel human MSN FERM structure (6TXQ) with a previous structure (1E5W). FERM subdomains of 6TXQ are coloured as in Fig. 1B, while 1E5W is coloured pink. (B) Top view of overlaid 6TXQ and 1E5W structures, with alignment over the F1 subdomain. The F3 subdomain in 6TXQ is rotated 12° along the axis of the long alpha helix, as measured by DynDom. (C) Overlay of individual subdomains from 6TXQ and 1E5W. Both F1 and F2 subdomains have small RMSD values and are closely aligned. In contrast, the F3 subdomain has a large RMSD value, with "open" and "closed" conformations observed for the previous (1E5W) and new (6TXQ) structures, respectively. RMSD=root mean square deviation.

### Figure S2

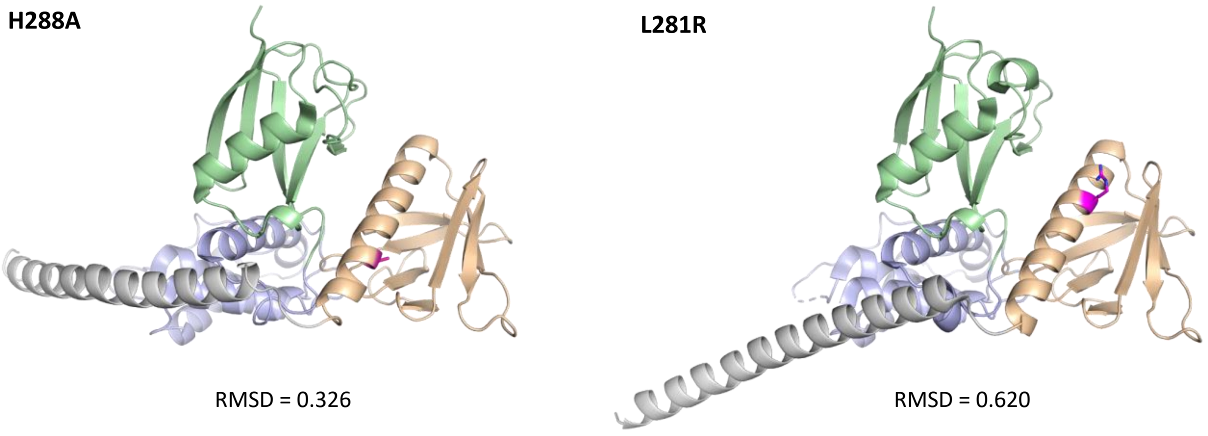


**Figure S2**: Structures of MSN FERM domain containing either H288A or L281R point mutations show minimal differences to wild-type FERM domains.

The crystal structures of mutant FERM domains of human MSN, containing the point mutations H288A (PDB: 8CIU) or L281R (PDB: 8CIT). Shown are cartoon models, with the FERM subdomains coloured the same way as in Fig. 1B. The mutated residues are highlighted in magenta. The RMSD values of mutant FERM domains, compared to wild-type (PDB: 6TXQ), are low, showing minimal differences in protein folding. In the case of MSN-L281R, the RMSD value is an average from three molecules in the asymmetric unit. RMSD=root mean square deviation.

### Figure S3

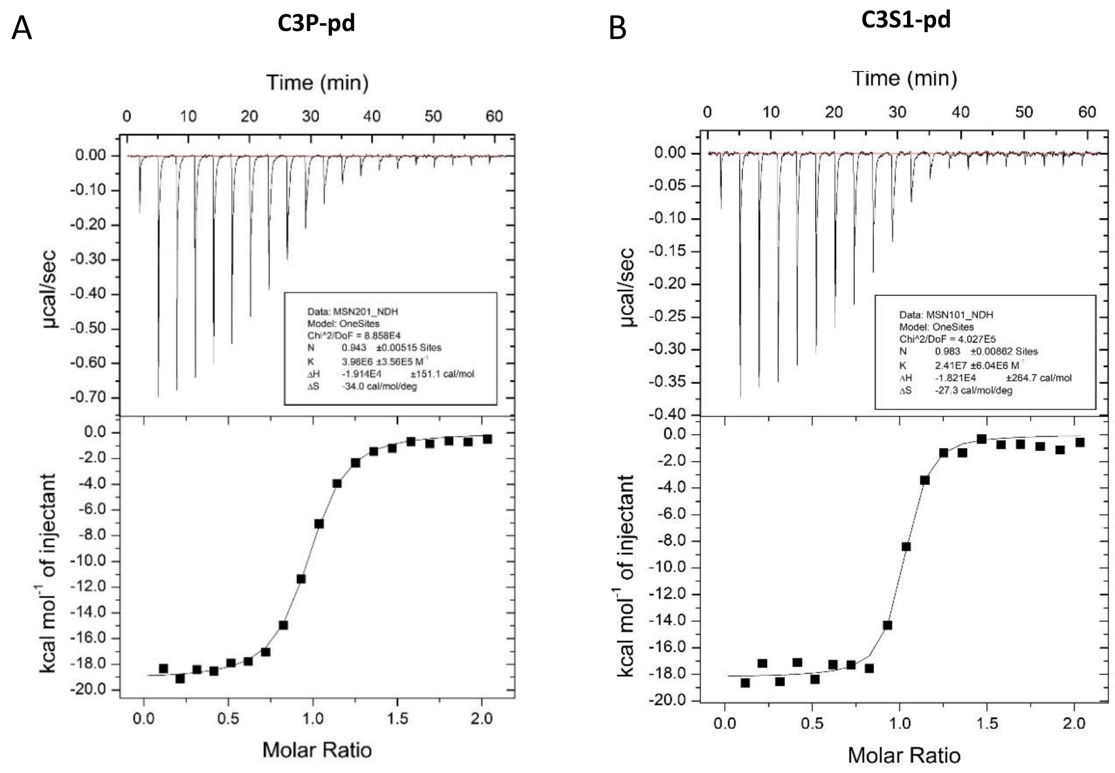


**Figure S3**: Binding kinetics of phage display peptides by ITC

(**A**, **B**) Example ITC isotherms of MSN FERM domain with peptide injections (upper) and integrated peak fit (lower) using a 1:1 binding model. (**A**) C3P-pd peptide. (**B**) C3S1-pd peptide.

**Table S1:** Crystallographic and refinement statistics

| | **Apo** | **CD44** | **C3P-pd** | **C3P-pd, C3S1-pd** | **L281R** | **H288A** |
|---|---|---|---|---|---|---|
| Crystallographic statistics[a] | | | | | | |
| Space group | $P4_32_12$ | $I4_1$ | $P2_1$ | P2 | $P2_1$ | $P4_32_12$ |
| Cell dimensions (Å, °) | 112.4, 112,4 61.1<br>90, 90, 90 | 120.0, 120.0, 64.6<br>90, 90, 90 | 45.5, 155.4, 63.6<br>90, 94.3, 90 | 79.7, 37.3, 81.5<br>90, 108.2, 90 | 72.7, 74.0, 115.1<br>90, 95.6, 90 | 117.1, 117.1, 62.7<br>90, 90, 90 |
| Resolution (Å) | 112.4 – 1.73<br>(1.76-1.73) | 84.8 – 2.20<br>(2.27 – 2.20) | 77.7 – 1.85<br>(1.89 – 1.85) | 77.5 – 1.42<br>(1.56 – 1.42) | 62.2 – 2.54<br>(3.07 – 2.54) | 55.3 – 2.39<br>(2.76 – 2.39) |
| $R_{merge}$ | 0.260 (16.039) | 0.166 (3.189) | 0.188 (2.137) | 0.136 (2.662) | 0.352 (1.268) | 0.195 (4.308) |
| $R_{pim}$ | 0.038 (3.121) | 0.048 (3.172) | 0.078 (1.089) | 0.029 (0.532) | 0.143 (0.520) | 0.028 (0.632) |
| $CC_{1/2}$ | 0.999 (0.377) | 0.999 (0.460) | 0.997 (0.347) | 0.999 (0.707) | 0.976 (0.591) | 1.000 (0.787) |
| Mean <I/σI> | 15.6 (0.7) | 11.1 (0.6) | 7.8 (0.9) | 13.2 (1.4) | 4.3 (1.5) | 21.6 (1.6) |
| Completeness (%)<br>Spherical<br>Ellipsoidal | <br>100.0 (99.8)<br>N/A | <br>99.7 (96.4)<br>N/A | <br>100.0 (99.6)<br>N/A | <br>59.7 (12.1)<br>92.9 (25.3) | <br>58.5 (22.3)<br>92.1 (74.9) | <br>68.6 (29.3)<br>91.7 (72.9) |
| Total reflections | 3,655,189<br>(113,752) | 555,227<br>(23,205) | 989,296<br>(42,198) | 1,156,637<br>(65,333) | 165,129<br>(27,222) | 619,298<br>(82,705) |
| Total unique reflections | 41,463 (2,210) | 23,383 (1,977) | 74,812 (4,621) | 51,601 (2,581) | 23,727 (3,949) | 12,176 (1,758) |
| Multiplicity | 88.2 (51.5) | 23.7 (11.7) | 13.2 (9.1) | 22.4 (25.3) | 7.0 (6.9) | 50.9 (47.0) |
| Refinement statistics | | | | | | |
| $R_{work}/R_{free}$ | 0.200/0.234 | 0.232/0.276 | 0.178/0.241 | 0.182/0.227 | 0.244/0.288 | 23.8/30.8 |
| RMSDs<br>Bond lengths (Å)<br>Bond angles (°) | <br>0.010<br>1.552 | <br>0.007<br>1.470 | <br>0.011<br>1.635 | <br>0.011<br>1.727 | <br>0.009<br>1.026 | <br>0.011<br>1.157 |
| Ramachandran statistics (%)<br>Favoured<br>Allowed<br>Outliers | <br>97.5<br>2.5<br>0 | <br>95.1<br>4.7<br>0.3 | <br>98.4<br>1.6<br>0 | <br>97.9<br>2.1<br>0 | <br>96.2<br>3.6<br>0.2 | <br>94.9<br>4.5<br>0.6 |
| Average B-factors<br>Protein<br>Ligand of interest<br>Other ligand<br>Water | <br>42.0<br>N/A<br>59.8<br>40.3 | <br>64.6<br>64.2<br>N/A<br>51.4 | <br>34.6<br>32.4<br>43.0<br>38.8 | <br>24.8<br>31.3<br>38.9<br>33.3 | <br>38.2<br>N/A<br>N/A<br>N/A | <br>73.0<br>N/A<br>N/A<br>42.6 |
| Number of Atoms<br>Protein<br>Ligand of interest<br>Other ligand<br>Water | <br>2714<br>N/A<br>4<br>164 | <br>2837<br>68<br>N/A<br>39 | <br>5793<br>210<br>89<br>844 | <br>2670<br>224<br>68<br>461 | <br>8259<br>N/A<br>N/A<br>N/A | <br>2702<br>N/A<br>N/A<br>86 |
| PDB Code | 6TXQ | 6TXS | 8CIR | 8CIS | 8CIT | 8CIU |

[a]outer shell statistics are given in brackets

**Table S2:** Structure activity relationship studies of compound 1

**Structure**

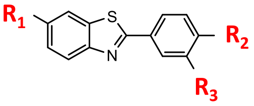


| Compound | $R_1$ | $R_2$ | $R_3$ | MSN TR-FRET ($IC_{50}$) | SYK TR-FRET ($IC_{50}$) |
|---|---|---|---|---|---|
| 1 | 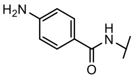 | 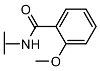 | H | 8.1 µM | ND |
| 1a | 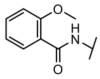 | 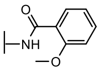 | H | 47.7 µM | 17.4 µM |
| 1b | 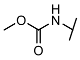 | 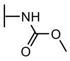 | H | 2.2 µM | 5.4 µM |
| 1c | H | 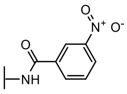 | H | >100 µM | >100 µM |
| 1d | H | 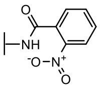 | H | >100 µM | >100 µM |
| 1e | H | 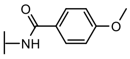 | H | >100 µM | >100 µM |
| 1f | H | 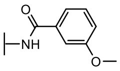 | H | >100 µM | >100 µM |
| 1g | H | 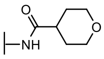 | H | >100 µM | >100 µM |
| 1h | H | 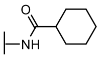 | H | >100 µM | >100 µM |
| 1i | H | H | 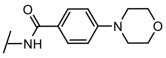 | >100 µM | >100 µM |
| 1j | 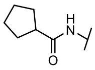 | 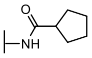 | H | >100 µM | >100 µM |
| 1k | 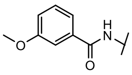 | 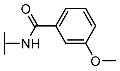 | H | >100 µM | >100 µM |
| 1l | H | H | 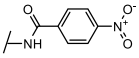 | >100 µM | >100 µM |
| 1m | 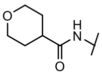 | 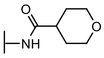 | H | >20 µM | >20 µM |
| 1n | 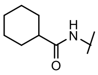 | 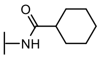 | H | >20 µM | >20 µM |
| 1o | H | 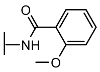 | H | >100 µM | >100 µM |
| 1p | 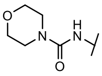 | 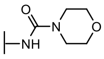 | H | >100 µM | >100 µM |

**Table S3:** Structure activity relationship studies of compound 2

| Compound | Structure<br>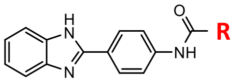<br>R | MSN TR-FRET ($IC_{50}$) | SYK TR-FRET ($IC_{50}$) |
|---|---|---|---|
| 2 | 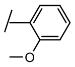 | 13.4 µM | 18.6 µM |
| 2a | 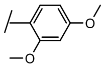 | 1.6 µM | >100 µM |
| 2b | 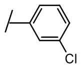 | 6.3 µM | >100 µM |
| 2c | 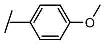 | 8.0µM | 14.0 µM |
| 2d | 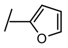 | 33.8 µM | 72.6 µM |
| 2e | 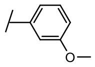 | >100 µM | >100 µM |
| 2f | 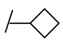 | >100 µM | >100 µM |
| 2g | 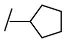 | >100 µM | >100 µM |
| 2h | 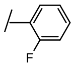 | >100 µM | >100 µM |
| 2i | 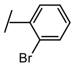 | >100 µM | >100 µM |
| 2j | 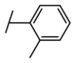 | >100 µM | >100 µM |
| 2k | 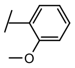 | >100 µM | >100 µM |
| 2l | 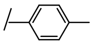 | >100 µM | >100 µM |

### Emory-Sage-SGC TREAT-AD Center Members

Ishita Ajith[1], Joel K. Annor-Gyamfi[2], Jeff Aube[2], Alison D. Axtman[2], Frances M. Bashore[2], Ranjita S. Betarbet[3], Juan Botas[4], William J. Bradshaw[1], Paul E. Brennan[1], Peter J. Brown[2], Robert R. Butler 3rd[5], Jacob L. Capener[2], Gregory W. Carter[6], Gregory A. Cary[6], Catherine Chen[4], Rachel Commander[3], Sabrina Daglish[2], Suzanne Doolen[7], Yuhong Du[3], Aled M. Edwards[8], Michelle E. Etoundi[4], Kevin J. Frankowski[2], Stephen V. Frye[2], Haian Fu[3], Opher Gileadi[1], Marta Glavatshikh[2], Jake Gockley[9], Katerina Gospodinova[1], Anna K. Greenwood[9], Peter A. Greer[10], Lea T. Grinberg[11], Shiva Guduru[2], Levon Halabelian[8], Crystal Han[5], Brian Hardy[2], Laura M. Heath[9], Stephanie Howell[2], Andrey A. Ivanov[3], Suman Jayadev[12], Vittorio L. Katis[1], Stephen Keegan[6], May Khanna[13], Dmitri Kireev[2], Carl LaFlamme[14], Karina Leal[9], Tom V. Lee[4], Tina M. Leisner[2], Allan I. Levey[3], Qianjin Li[3], David Li-Kroeger[4], Zhandong Liu[4], Benjamin A. Logsdon[9], Frank M. Longo[5], Lara M. Mangravite[9], Peter S. McPherson[14], Richard M. Nwakamma[3], Felix O. Nwogbo[2], Carolyn A. Paisie[6], Arti Parihar[12], Kenneth H. Pearce[2], Kun Qian[3], Min Qui[3], Stacey J Sukoff Rizzo[7], Karolina A. Rygiel[1], Julie Schumacher[5], David D. Scott[15], Nicholas T. Seyfried[3], Joshua M. Shulman[4], Ben Siciliano[3], Arunima Sikdar[2], Nathaniel Smith[4], Michael Stashko[2], Judith A. Tello Vega[15], Dilipkumar Uredi[2], Dongxue Wang[3], Jianjun Wang[3], Xiaodong Wang[2], Zhexing Wen[3], Jesse C. Wiley[9], Alexander Wilkes[1], Charles A. Williams[12], Timothy M. Willson[2], Aliza Wingo[3], Thomas S. Wingo[3], Novak Yang[3], Jessica E. Young[12], Miao Yu[6], Elizabeth L. Zoeller[3]

[1]University of Oxford, Oxford, OX3 7FZ, UK
[2]University of North Carolina, Chapel Hill, NC 27599, USA
[3]Emory University School of Medicine, Atlanta, GA 30322, USA
[4]Baylor College of Medicine, Houston, TX 77030, USA
[5]Stanford University School of Medicine, Stanford, CA, 94305, USA
[6]The Jackson Laboratory, Bar Harbor, ME 04609, USA
[7]University of Pittsburgh School of Medicine, Pittsburgh, PA 15219, USA
[8]University of Toronto, Toronto, ON M5G 1L7, Canada
[9]Sage Bionetworks, Seattle, WA, 98121, USA
[10]Queen's University, Kingston, Ontario, ON K7L 3N6, Canada
[11]University of California, San Francisco, San Francisco, CA 94143, USA
[12]University of Washington, Seattle, WA 98109, USA
[13]New York University, New York, NY 10010, NY, USA
[14]McGill University, Montreal, QC H3A 2B4, Canada
[15]University of Arizona, Tucson, AZ 85724, USA

### Chemical Synthesis: 1H NMR and LCMS analysis

#### General Information

Reagents were obtained from vetted commercial suppliers and used without purification. Temperatures are reported in degrees celsius (°C); solvent was removed under reduced pressure using a rotary evaporator; and reaction progress was tracked via thin layer chromatography. The following abbreviations are used in experimental procedures: mmol (millimoles), µmol (micromoles), mg (milligrams), equiv (equivalent(s)), rt (room temperature), mins (minutes), and h (hours). $^{1}$H NMR and additional microanalytical data were collected for final compounds to confirm their identity and evaluate their purity. $^{1}$H NMR spectra were obtained in DMSO-$d_6$ and recorded using Bruker spectrometers. Magnet strength is included in the line listing for each compound. Peak positions are reported in parts per million (ppm) and calibrated versus the shift of DMSO-$d_6$ and coupling constants (*J* values) in hertz (Hz). Peak multiplicities are included as follows: singlet (s), doublet (d), doublet of doublets/triplets (dd/dt), triplet (t), triplet of doublets (td), doublet of doublet of doublets (ddd), and multiplet (m). Purity was determined using high-performance liquid chromatography (HPLC) and all final compounds demonstrated >95% purity.

*4-amino-N-(4-(6-(4-aminobenzamido)benzo[d]thiazol-2-yl)phenyl)benzamide bis(2,2,2-trifluoroacetate)* ***(1)***

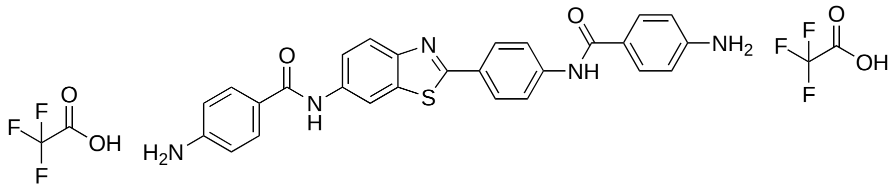


#### $^{1}$H NMR

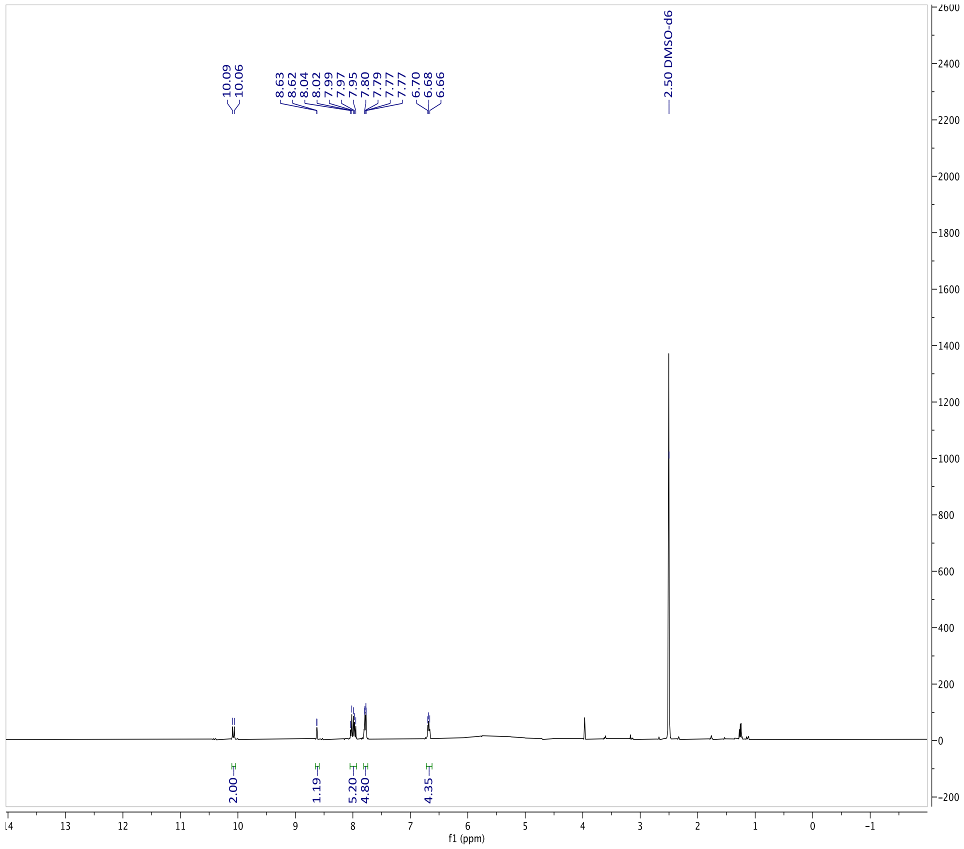

### LCMS

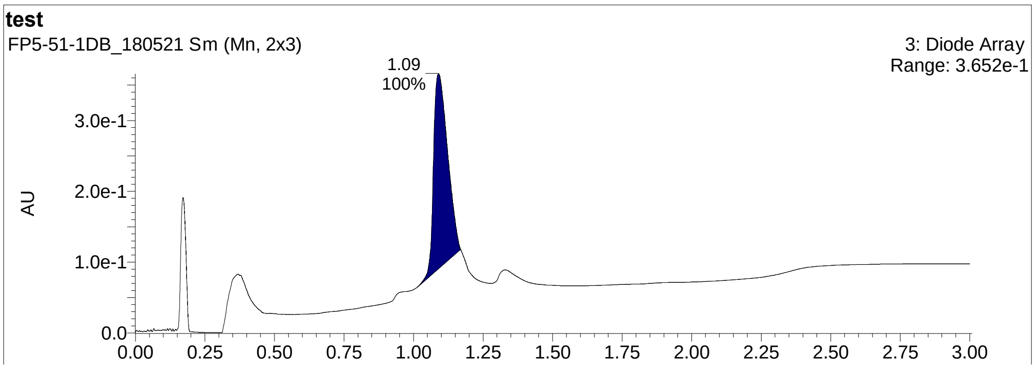
test
FP5-51-1DB_180521 Sm (Mn, 2x3)
3: Diode Array
Range: 3.652e-1
1.09
100%
AU
3.0e-1
2.0e-1
1.0e-1
0.0
0.00
0.25
0.50
0.75
1.00
1.25
1.50
1.75
2.00
2.25
2.50
2.75
3.00

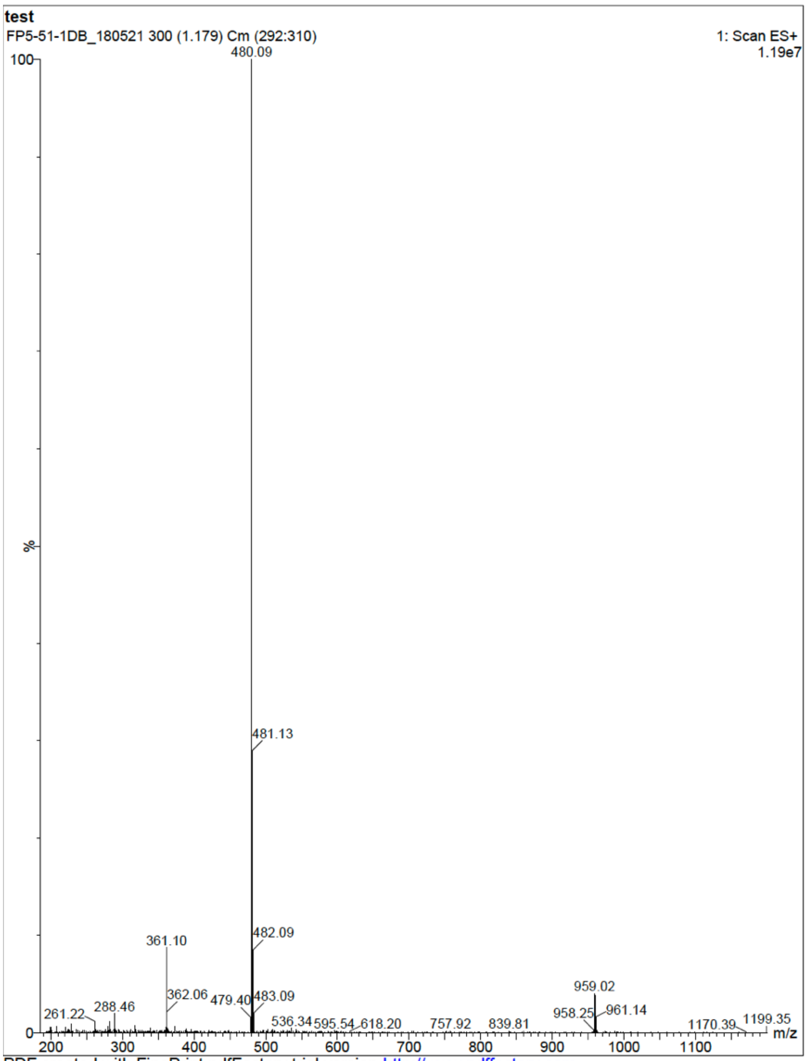
test
FP5-51-1DB_180521 300 (1.179) Cm (292:310)
1: Scan ES+
1.19e7
480.09
100
%
0
481.13
482.09
361.10
959.02
362.06
479.40
483.09
261.22
288.46
536.34
595.54
618.20
757.92
839.81
958.25
961.14
1170.39
1199.35
m/z
200
300
400
500
600
700
800
900
1000
1100

*N-(4-(1H-benzo[d]imidazol-2-yl)phenyl)-2-methoxybenzamide (**2**)*

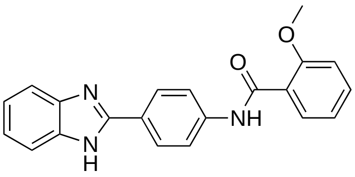

**[1]H NMR**

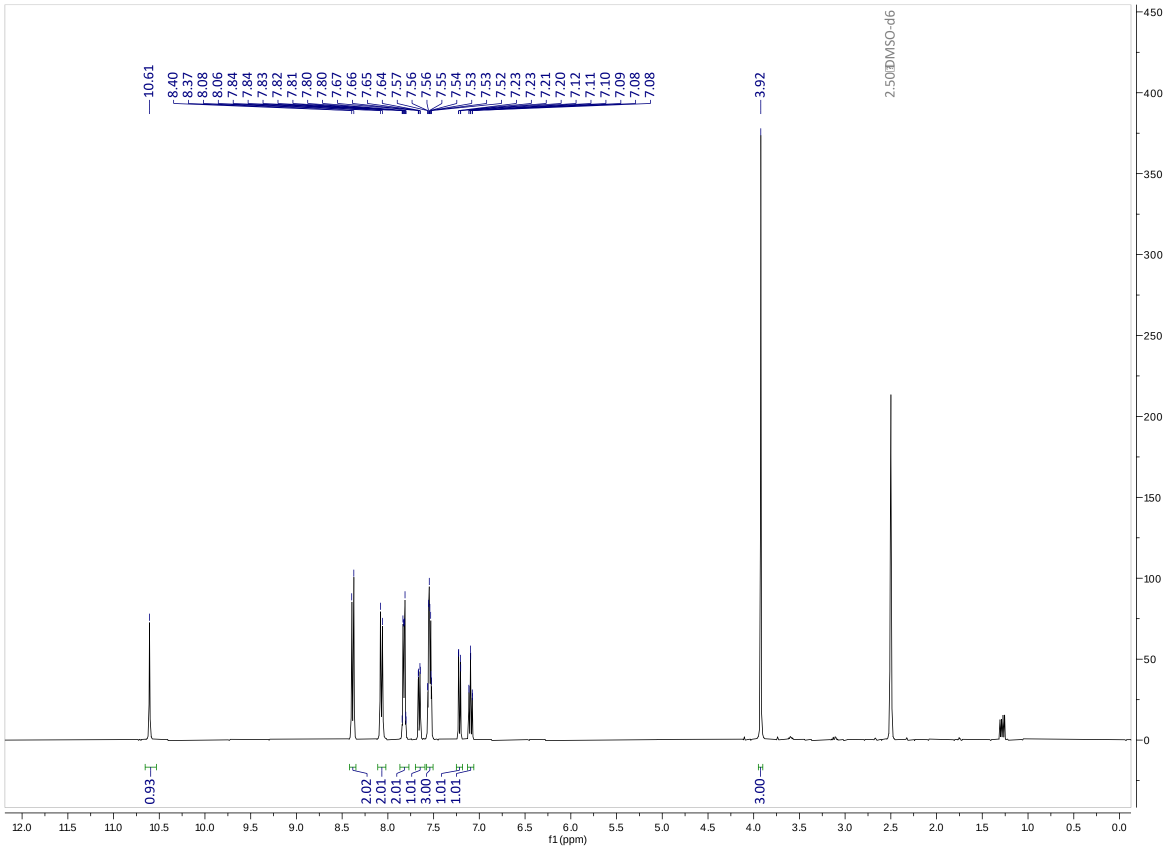

**LCMS**

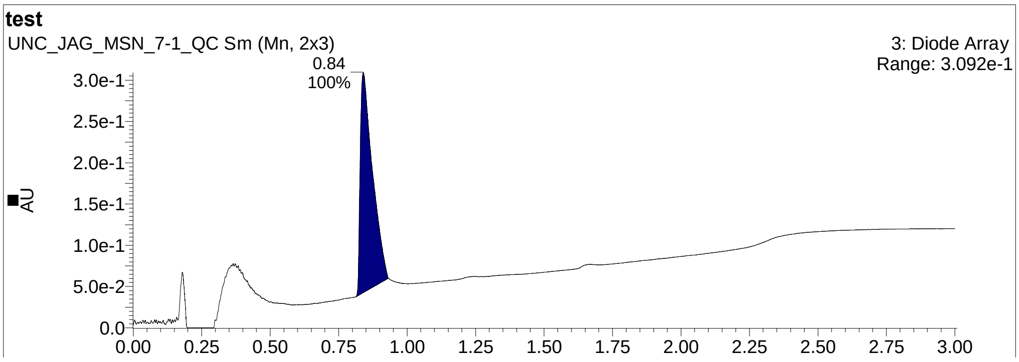

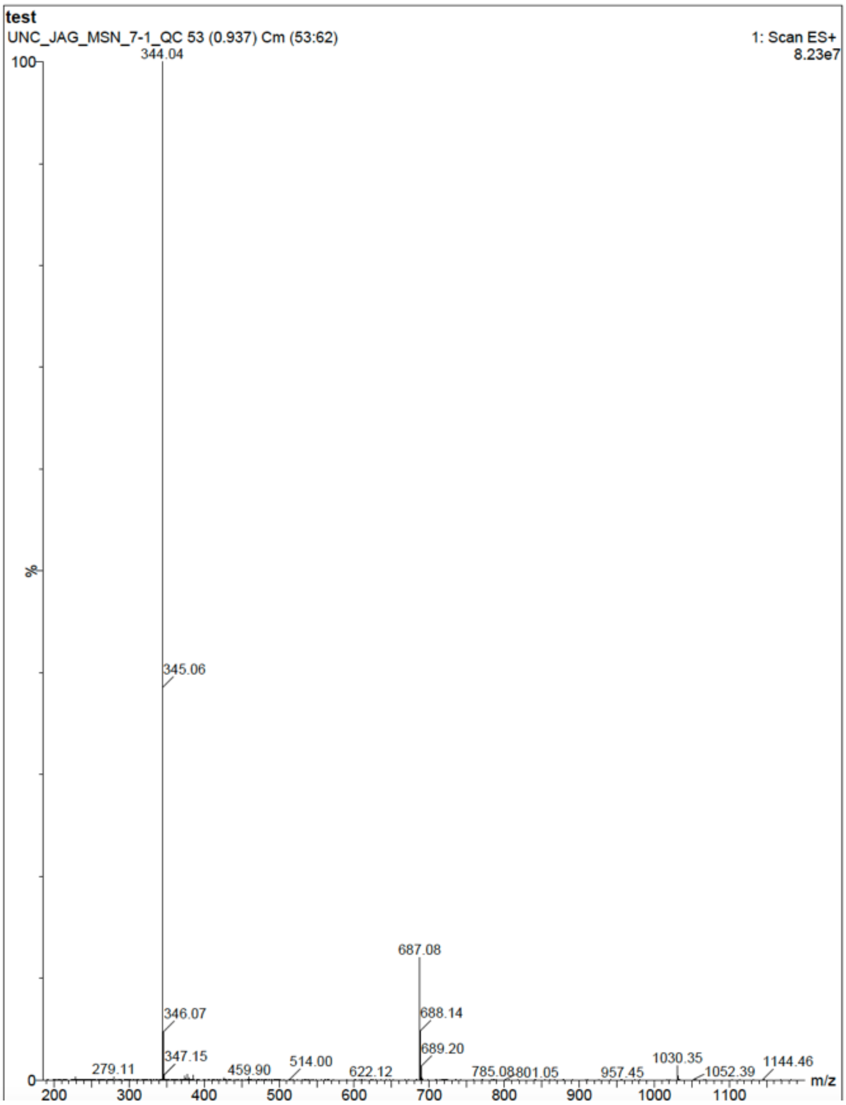
test
UNC_JAG_MSN_7-1_QC 53 (0.937) Cm (53:62)
1: Scan ES+
8.23e7
344.04
345.06
687.08
346.07
688.14
347.15
689.20
279.11
459.90
514.00
622.12
785.08
801.05
957.45
1030.35
1052.39
1144.46
%
m/z

*N-(4-(1H-benzo[d]imidazol-2-yl)phenyl)-2,4-dimethoxybenzamide (**2a**)*

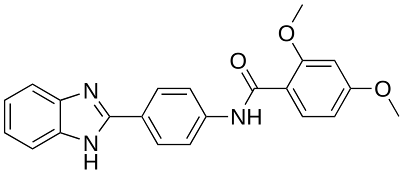


### [1]H NMR

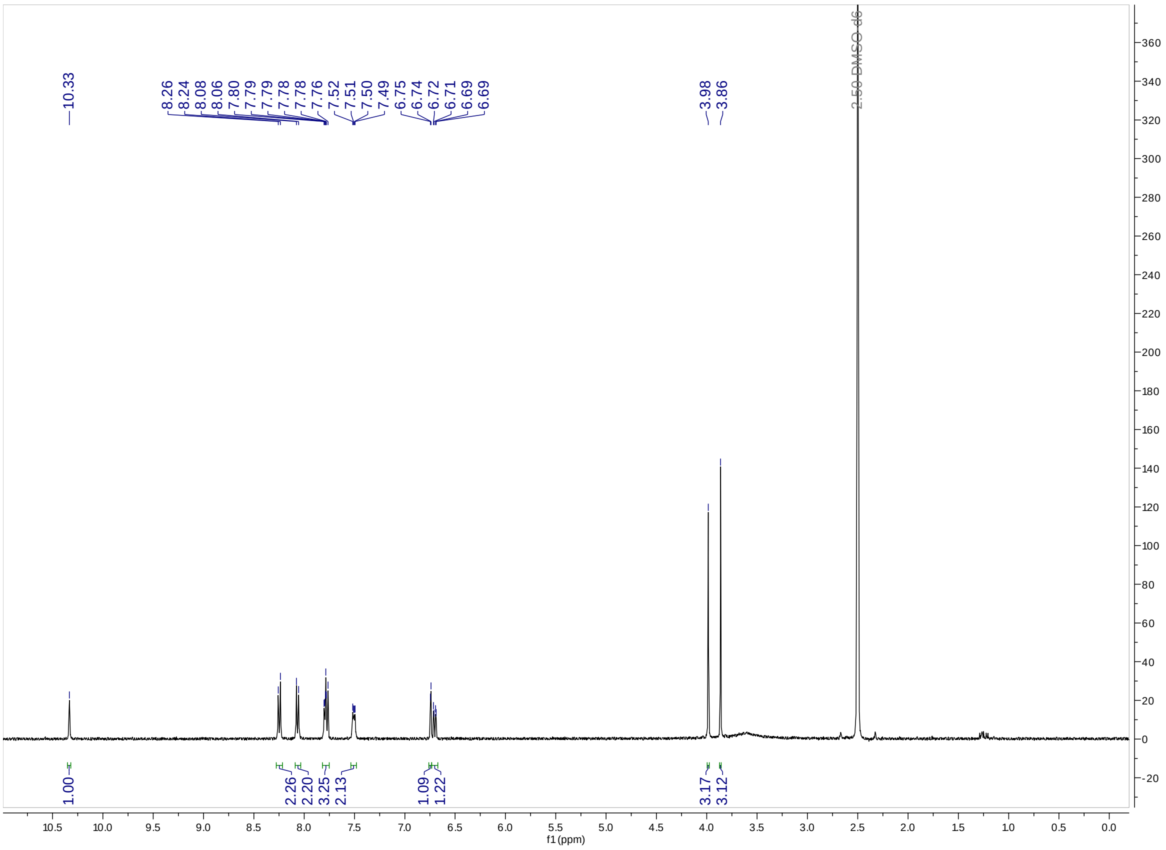


### LCMS

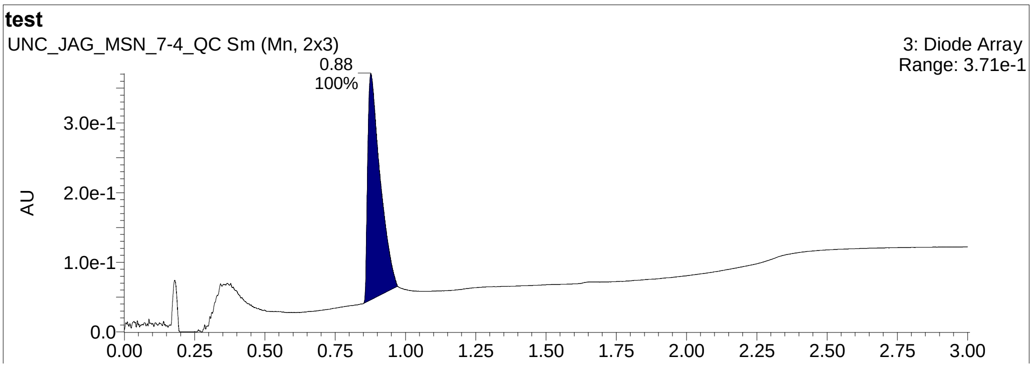

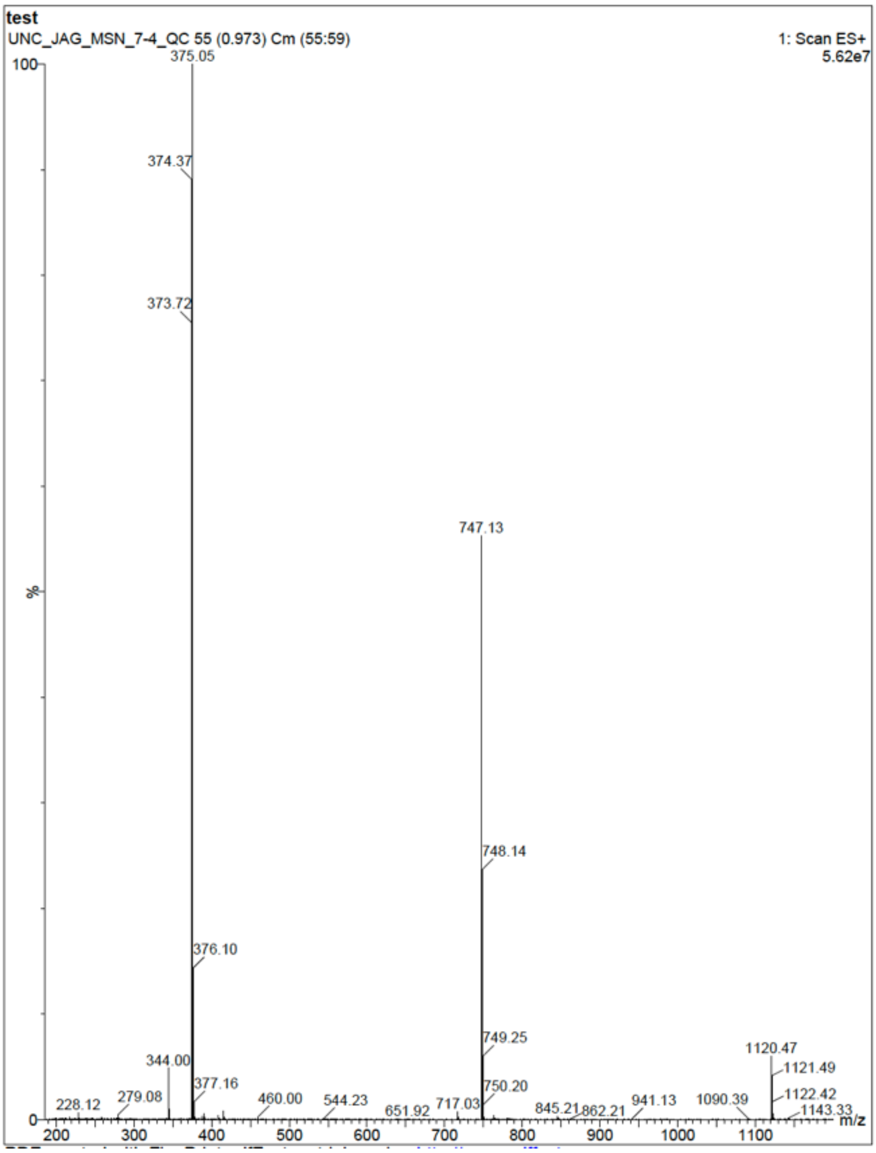
test
UNC_JAG_MSN_7-4_QC 55 (0.973) Cm (55:59)
1: Scan ES+
5.62e7
375.05
374.37
373.72
747.13
748.14
376.10
749.25
344.00
377.16
750.20
1120.47
1121.49
1122.42
1143.33
228.12
279.08
460.00
544.23
651.92
717.03
845.21
862.21
941.13
1090.39
%
m/z

*N-(4-(1H-benzo[d]imidazol-2-yl)phenyl)-3-chlorobenzamide (**2b**)*

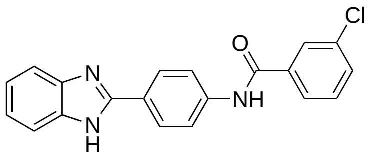


**[1]H NMR**

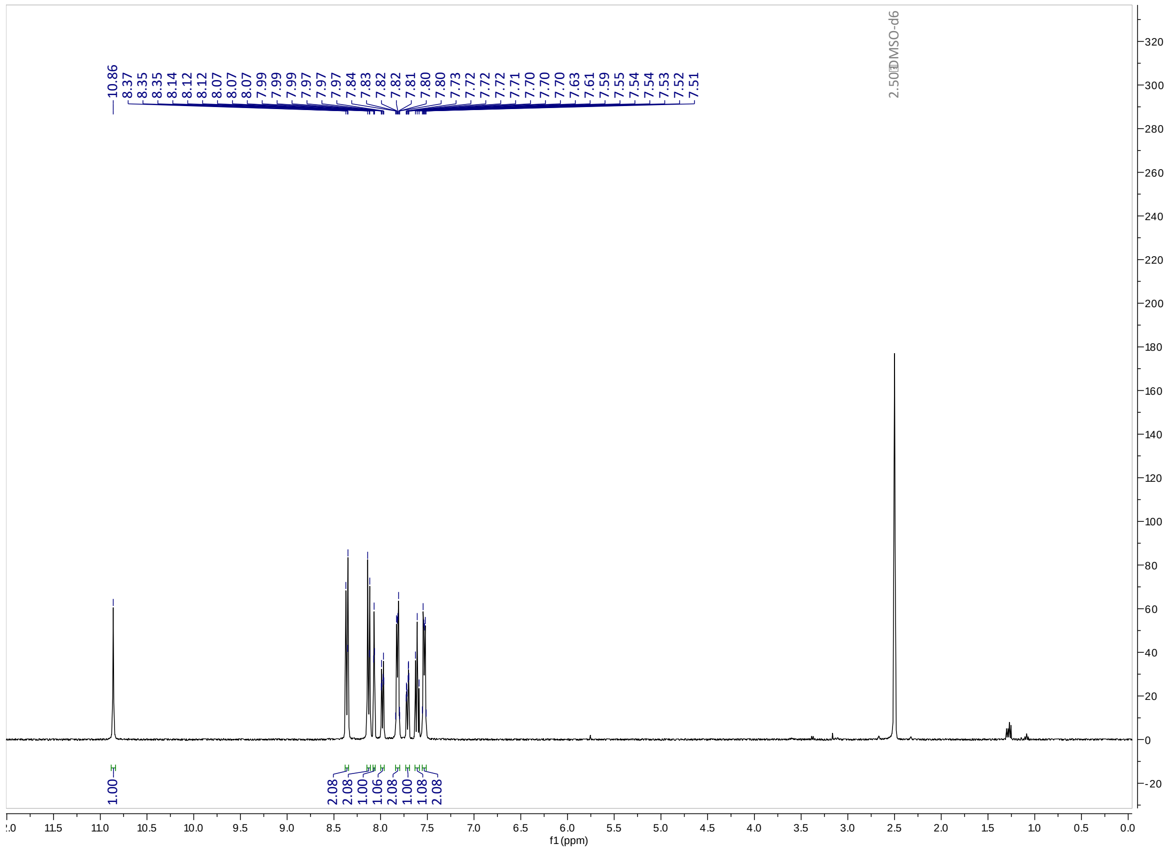


**LCMS**

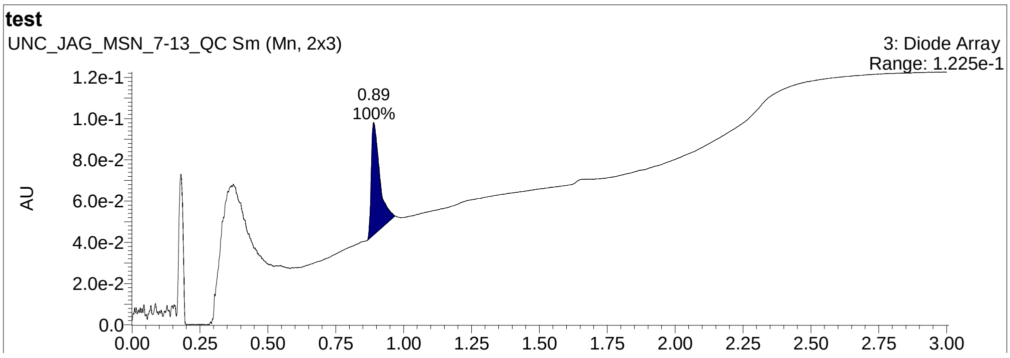

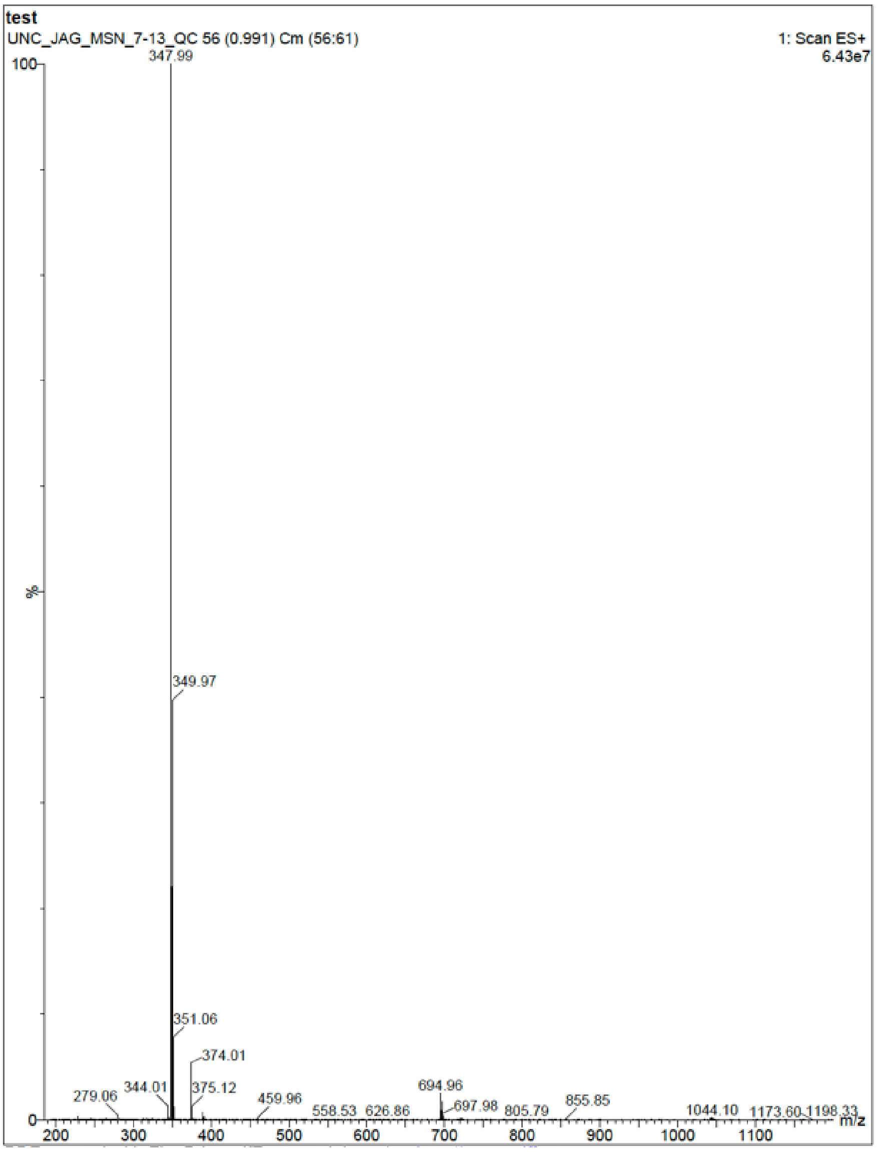
test
UNC_JAG_MSN_7-13_QC 56 (0.991) Cm (56:61)
1: Scan ES+
6.43e7
347.99
349.97
351.06
374.01
375.12
279.06
344.01
459.96
558.53
626.86
694.96
697.98
805.79
855.85
1044.10
1173.60
1198.33
%
m/z
